# Stop codon readthrough of a POU transcription factor regulates steroidogenesis and developmental transitions

**DOI:** 10.1101/2020.05.30.125401

**Authors:** Yunpo Zhao, Bo Gustav Lindberg, Shiva Seyedoleslami Esfahani, Xiongzhuo Tang, Stefano Piazza, Ylva Engström

**Affiliations:** Department of Molecular Biosciences, The Wenner-Gren Institute, Stockholm University, SE-106 91 Stockholm, Sweden

## Abstract

Translational stop codon readthrough generates C-terminally extended protein isoforms. While evidence mounts of readthrough as a global phenomenon, proofs of its functional consequences are scarce. We show that readthrough of the mRNA for the *Drosophila* POU/Oct transcription factor Drifter occurs at a high rate and in a spatiotemporal manner *in vivo*, reaching above 50% in the prothoracic gland. Phylogenetic analyses suggested that readthrough of *drifter* is conserved among Dipterans, with C-terminal extensions harboring intrinsically disordered regions, and amino acids streches implied in transcriptional activation. The C-terminally extended Drifter isoform is required for maintaining normal levels of the growth hormone ecdysone through regulation of its biosynthetic genes, acting in synergy with the transcription factor Molting defective. A 14-bp deletion that abolished readthrough, caused prolonged larval development and delayed metamorphosis. This study provides a striking example of alternative genetic decoding that feeds into the progression from one life cycle stage to another.

## Introduction

From the last decades of genome and metagenome sequencing projects it has become apparent that the genetic code is non-universal, as a repertoire of alternative genetic decoding exists (Baranov et al., 2015; Rodnina et al., 2019). The added decoding plasticity increases the protein repertoire without expanding the genome, which could benefit organisms undergoing spatiotemporal alterations of the proteome in response to certain cues.

Translation of mRNA by the ribosome continues until a stop codon (UAA, UAG, or UGA) is reached, which allows release factors to recognize the stop codon and mediate termination. Normally, the error rate of termination is less than 0.1%. In some cases, however, ribosomes interpret the stop codon as a sense codon, resulting in stop codon readthrough (SCR). Initially characterized as an evolved common strategy of viruses to expand the proteome (Beier et al., 1984; Pelham, 1978; Weiner and Weber, 1971), SCR has recently been documented to occur in yeast, fungi, plants, insects, nematodes and mammals, where it seems to act as gene regulatory mechanism for the synthesis of protein isoforms with extended C-termini. The added domains can provide signals for protein sorting, localization, stabilization/destabilization and other functional domains (Baranov et al., 2015).

In eukaryotic cells, the tRNA-shaped eukaryotic release factor eRF1 recognizes the three stop codons and facilitates the release of nascent polypeptide chains from the peptidyl-tRNA positioned at the ribosome P-site (Hellen, 2018). If the interaction between eRF1 and mRNA is not efficient enough, near-cognate tRNAs (nc-tRNAs) are able to decode the stop codons as sense codons, leading to SCR. The identity of the stop codon contributes to the relative termination fidelity, with UGA having the highest SCR potential, followed by UAG and UAA (Cridge et al., 2018).

The base immediately 3’ of the stop codon also affects the readthrough, e.g. the level of UGA-C readthrough is higher than that of UGA-N (Beznoskova et al., 2016). In addition, RNA stem-loop structures are enriched in the vicinity of potentially leaky stop codons, and is both sufficient and necessary for readthrough of the *headcase (hdc)* gene in *Drosophila* (Steneberg and Samakovlis, 2001). In a few cases, RNA binding proteins and miRNAs have been found to control the rate of SCR. For example, heterogeneous nuclear ribonucleoprotein (hnRNP) A2/B1 was shown to bind to a *cis*-acting element in *VEGF-A* 3’ untranslated region (UTR) and promote SCR (Eswarappa et al., 2014). Translational readthrough of the mammalan *AGO1* gene, encoding the Argonaute 1 (Ago1) protein, was recently found to be positively regulated by the let-7a miRNA upon binding 3’ of the canonical stop codon (Singh et al., 2019).

Several whole genome approaches have been used to identify genes undergoing SCR, such as ribosome profiling, phylogenetic analyses of codon substitution frequencies (PhyloCSF) and *in silico* identification of genes with specific stop codon contexts that are more prone to SCR (Schueren and Thoms, 2016). For example, 57 human genes were identified with a favourable stop codon context and six of these were experimentally verified (Stiebler et al., 2014). A recent annotation of SCR in nine vertebrate model organisms resulted in 13 genes exhibiting phylogenetically conserved C-terminal extensions, in total resulting in 94 SCR isoforms (Rajput et al., 2019). The most pervasive whole genome analyses of SCR have been carried out in insects, taking advantage of the complete genome sequences of numerous *Drosophila* and *Anopheles* species. Comparative genome analysis of 12 *Drosophila* species initially predicted that 149 genes undergo SCR (Stark et al., 2007). In follow-up studies of 20 *Drosophila* and 21 *Anopheles* species, SCR was predicted for a total of 333 *Drosophila* and 353 *Anopheles* genes (Jungreis et al., 2016; Jungreis et al., 2011). Deep sequencing of ribosome-protected mRNA fragments (a.k.a. ribosome profiling) have provided genome-wide experimental validation of SCR in *Drosophila* (Dunn et al., 2013) and in mammalian cells (Wangen and Green, 2020).

The functional importance of SCR has, however, only been sparsely investigated. Early experimental studies identified the *Drosophila* genes for *Synapsin, kelch* and *hdc* to produce alternative protein products through SCR (Klagges et al., 1996; Robinson and Cooley, 1997; Steneberg et al., 1998). SCR of *hdc* mRNA was shown to contribute to the regulation of tracheal development, providing one of the first evidences of an essential role of C-terminally extended proteins in *Drosophila* (Steneberg and Samakovlis, 2001). Except these cases, a clear understanding of the biological context and functional roles *in vivo* of SCR is essentially lacking.

A relatively large number of SCR candidate genes are transcription factors, out of which several are involved in nervous system development (Pancsa et al., 2016). An interesting gene in this respect is *Drosophila drifter (dfr)/ventral veins lacking (vvl)*, (from hereon referred to as *dfr),* which is predicted to encode an unusually long C-terminal extension upon readthrough (Jungreis et al., 2016; Jungreis et al., 2011). Dfr is a member of the POU/Oct domain transcription factor family, including well-known regulators of embryonic and neural development, stem cell pluripotency, immunity and cancer. Dfr plays profound roles during all stages of *Drosophila* development, such as regulation of embryonic brain and nervous system development, tracheogenesis, and adult epithelial immunity (Anderson et al., 1995; Certel and Thor, 2004; de Celis et al., 1995; Junell et al., 2010). Its mammalian orthologs, POU3F1-POU3F4 regulate embryogenesis, neurogenesis and neuronal differentiation and are referred to as the POU-class III of neural transcription factors.

POU/Oct proteins also control developmental transitions, such as POU1F1/Pit1, which in mammals regulates expression of several genes involved in pituitary development, expression of growth hormone and prolactin, and progression of puberty. Similarly, it has been shown that *dfr* controls metamorphosis in insects by controlling the synthesis and release of steroid hormones from the prothoracic gland (PG) (Cheng et al., 2014; Danielsen et al., 2014), an endocrine organ with analogous functions to the mammalian pituitary gland.

In the present study, we show that the kinetic profile of ecdysteroid biosynthesis, which acts as a timekeeper and coordinator of insect metamorphosis, requires the activity of an SCR-derived, C-terminally extended isoform of Dfr, thus demonstrating a critical role of SCR for developmental transitions. The evolutionary conservation of SCR in metazoans implies that it may serve as a general regulatory mechanism, playing more profound roles in cellular and organismal processes than previously anticipated.

## Results

### Translational stop codon readthrough of *dfr* mRNA

Phylogenetic analyses of codon substitution frequencies (PhyloCSF) and *in silico* identification of genes with specific stop codon contexts have pointed out *dfr* as a strong candidate for SCR (Stark et al., 2007). Similar to its orthologs, including human POU3F1-4, the *dfr* locus is intronless with a long 3’ UTR (Figure 1A). The first open reading frame (ORF) produces a 45.9 kDa protein (from hereon referred to as Dfr-S, with S depicting the short form), whereas SCR into ORF2 would extend it by 286 amino acids to 76.8 kDa (referred to as Dfr-L, with L indicating the long form; Figure 1A). In addition, the next two downstream stop codons may also be subject to SCR, producing additional 78.1 and 79.9 kDa isoforms respectively (Jungreis et al., 2011). We hypothesized that such a long, evolutionarily conserved, C-terminal extension would provide additional, or altogether different, properties to the protein. The extent of SCR was analyzed using two different antibodies; one directed against the common N-terminal ORF1 (anti-Dfr-N), recognizing both Dfr-S and Dfr-L isoforms, and another directed against the C-terminal ORF2 (anti-Dfr-C), specific for Dfr-L (Figure 1A). In adult extracts from several strains of *Drosophila*, bands corresponding to the predicted molecular weights of both Dfr-S and Dfr-L were produced (Figure 1-figure supplement 1A and 1B). In addition, a 77 kDa band (Dfr-L) was expressed from *dfr* cDNA in an *in vitro*-coupled transcription/translation system (Figure 1-figure supplement 1C). In embryos of mixed stages, however, we and others (Anderson et al., 1995) only detected the Dfr-S isoform (Figure 1-figure supplement 1D), indicating that *dfr* is not subject to prominent SCR during embryogenesis. Lack of alternative splicing or potential RNA editing proximal to the stop codon was experimentally confirmed by DNA sequencing of a reverse-transcribed mRNA (Figure 1-figure supplement 1E and F). Taken together, these results indicate that SCR of *dfr* occurs both *in vivo* and *in vitro*.

**Figure 1.**
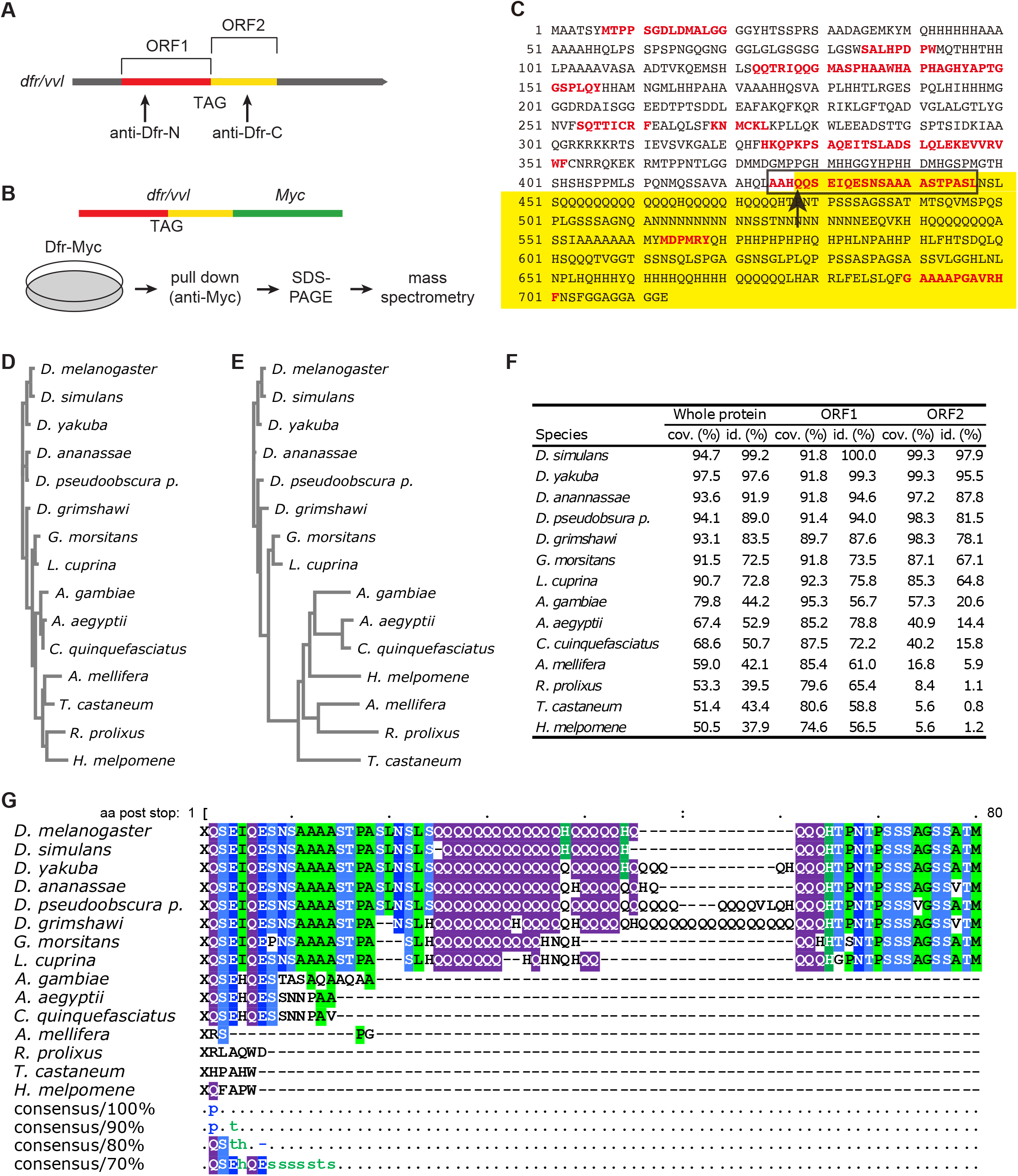
Translational readthrough of *dfr* mRNA produces two alternative Dfr isoforms. (**A**) Schematic representation of the intron-less *dfr/vvl* gene. The *dfr* open reading frame 1 (ORF1, 427 amino acids, red), ending at the first UAG stop codon, is followed directly by a second ORF2 (286 amino acids, yellow), flanked by 5’ and 3’ untranslated regions (gray). Translational readthrough of the UAG leads to uninterrupted translation and produces Dfr-L, with a C-terminal extension. Two independent Dfr antibodies were used in this study, recognizing the common N-terminal part of Dfr (anti-Dfr-N) and the Dfr-L-specific C-terminal extension (ORF2) (anti-Dfr-C) respectively. (**B**) Schematic illustration of expression and isolation of Dfr-L followed by mass spectrometry. *dfr* cDNA was ligated in the 3’ end of the predicted ORF2, to create an in-frame fusion with 6 x *Myc* tag and expressed in *Drosophila* cell culture. Expression of the Dfr-L-Myc fusion protein can only occur as a result of SCR, and it was pulled-down with a Myc antibody followed by separation with electrophoresis, digestion with chymotrypsin and analyzed by nano Liquid Chromatography-Mass Spectrometry (nLC-MS). (**C**) Results from the nLC-MS showing the Dfr-L amino acid sequence, with Dfr C-terminal extension highlighted in yellow. Nine peptides (bold red) matched the predicted Dfr-L sequence. Two of these peptides are located within the Dfr C-terminal extension and one peptide (boxed sequence) encompasses the UAG stop codon, which is decoded as an X in sequence data bases (Jungreis et al., 2011). The nLC-MS revealed that the incorporated amino acid is a glutamine (Q, arrow). (**D-G**) Phylogenetic comparison of plausible amino acid sequences resulting from translation of the ORF downstream of the first stop codon (ORF2). ORF1 (D) was analysed in conjuction with ORF2 (E) to compare the degree of conservation. **(D-E)** Phylogenetic trees, constructed using the Neighbour-joining method, depict real branch distances between ORFs of denoted species. Note that non-dipteran species included in the analysis had a comparatively short ORF2. **(F)** Summary of the percent coverage and identity of ORF1, 2, or both, among selected species when compared to *D. melanogaster* Dfr. **(G)** MView alignment of the first 80 amino acids downstream of the stop codon (X) from respective species.

The first stop codon of *dfr* mRNA is a UAG triplet (Figure 1A), which has an intermediate relative potential for readthrough (UGA>UAG>UAA)(Cridge et al., 2018). The frequency of SCR of *dfr* mRNA may be positively influenced by the presence of a cytosine immediately 3’ of the stop codon (UAG-C) and a predicted RNA:RNA stem loop structure immediately down-stream of the stop codon (Jungreis et al., 2016). To experimentally verify SCR of *dfr* mRNA and to determine the amino acid decoded from the UAG stop codon, mass spectrometry analysis was applied (Figure 1 B). This resulted in nine peptides with sequences aligning to Dfr. Importantly, two peptides matched within the C-terminal extension and one encompassed the first in-frame UAG stop codon (Figure 1C). This provides experimental evidence of *dfr* mRNA undergoing SCR. The UAG stop codon was translated into glutamine (Figure 1C) and no other amino acid incorporations were detected at this site. This indicates that the UAG codon was interpreted as a CAG codon, as AAG and GAG would be translated into lysine and glutamic acid respectively. We conclude that the UAG stop codon in *dfr* mRNA can be used as a template for tRNA^gln^ base pairing and incorporation of glutamine.

### Stop codon readthrough of *dfr* appears evolutionarily conserved in Diptera

To investigate the evolutionary conservation of the Dfr C-terminal extension, we performed a phylogenetic analysis using amino acid sequences from ORF1 and ORF2 independently (Figure 1D-G). In common for both, the resulting trees displayed similar patterns of divergence, although ORF2 sequences revealed larger relative distanses (Figure 1D-E). A distinct separation between flies and other species was evident when ORF2 sequences were compared, whereas all shared at least 56 % identity in ORF1 to *Drosophila melanogaster*. This suggests that ORF2 has undergone faster divergence than ORF1. Within more closely related dipteran species including *Drosophila, Lucilia* and *Glossina*, ORF1 and ORF2 were conserved to a fairly similar degree (73.5-100 % and 64.8-97.9 % identity, respectively), suggesting that the functional role of the extended form is also preserved. Low conservation of the extended ORF2 was found among selected mosquitoes (40.2-57.3% sequence coverage; 14.4-20.6% identity). Interestingly, *dfr* SCR has been proposed to occur in the malaria mosquito *Anopheles gambiae* as well (Jungreis et al., 2016), despite the apparent unrelatedness of ORF2 compared to *Drosophila* (Figure 1G). We noted, however, that sequences proximal to the stop codon was well conserved (Figure 1G).

### Spatiotemporal regulation of *dfr* stop codon readthrough

We next analyzed the relative expression levels of Dfr-S and Dfr-L isoforms in different tissues and stages of development. Immunostaining using anti-Dfr-N and Dfr-C antibodies revealed that both Dfr isoforms are predominantly nuclear, indicating that SCR did not change the subcellular localization of Dfr (Figure 2A-I). The PG of all three larval instars stained intensively with both antibodies, as well as the ring gland in late stage embryos (Figure 2A, D, G and Figure 2-figure supplement 1F and I), indicating prominent SCR. This was confirmed in extracts of brain/ring gland complexes (BRGCs) (Figure 2J) with a relative abundance of Dfr-L of 47% in females and 53% in males, demonstrating a very high degree of SCR. It can be noted that both isoforms of Dfr migrated with slower mobility in larval compared to adult extracts, although bands with this migration pattern were also observed in adults tissue extracts (Figure 2 J, K and Figure 1-figure supplement 1A and 1B), possibly reflecting post-translational modifications. A developmental profile of BRGCs showed that SCR was more frequent in early L3 larvae and then declined (Figure 2K-L). A moderate level of *dfr* SCR was also evident in larval trachea (Figure 2B, E), brain and CNS of L2 larvae (Figure 2G), in adult fat body cells and oenocytes (Figure 2H and I), and in several other larval and adult tissues (Figure 2-figure supplement 1A-I). In contrast, Dfr-L was barely detectable in the male ejaculatory duct (Figure 2F), adult trachea and ureter cells, in which anti-Dfr-N staining was prominent (Figure 2-figure supplement 2I and (Junell et al., 2010). Thus, these tissues primarily expresses the Dfr-S isoform and seem resilient to *dfr* SCR. This indicates that SCR is a highly regulated process, ranging from 50% in tissues like the larval PG, to tissues with high *dfr* gene expression without prominent SCR, such as the male ejaculatory duct. This underscores that the rate of *dfr* SCR is not simply the result of leaky translational termination. We conclude that *dfr* undergoes SCR in a spatiotemporal manner, suggesting that it is programmed as part of a gene regulatory program.

**Figure 2.**
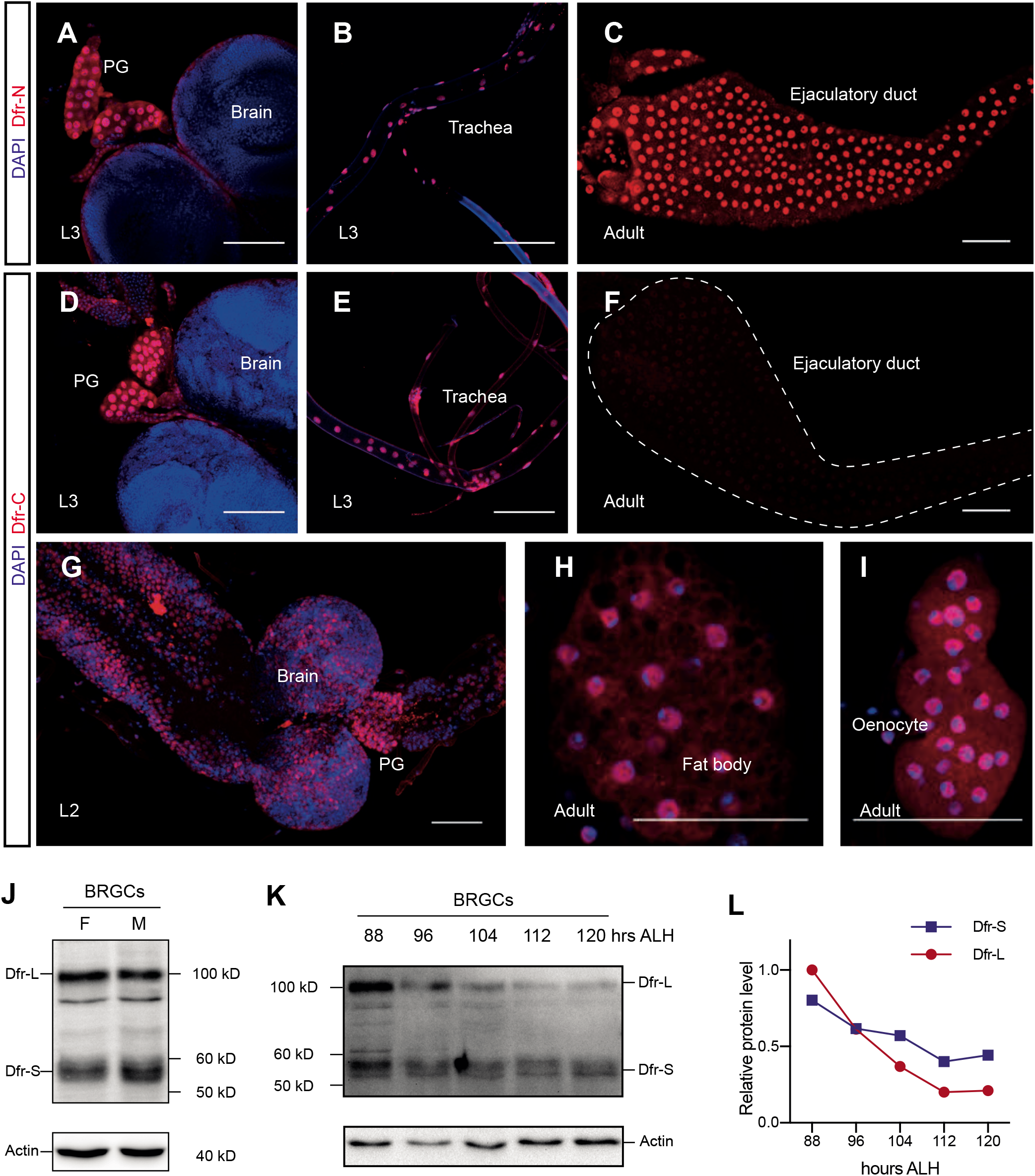
The relative frequency of *dfr* stop codon readthrough varies between tissues and development stages. (**A-I**) Confocal images of immunostaining, using the Dfr-N antibody recognizing both forms of Dfr (A-C), or the Dfr-C antibody specific for the C-terminal extension (D-I) of larval BRGCs (A, C, and G), larval trachea (B and E), adult male ejaculatory duct (C and F), adult fat body cells (H) and oenocytes (I). The Dfr-L isoform is prominently expressed in nuclei of the prothoracic gland (PG) and larval trachea, while it is barely detectable in the male ejaculatory duct (F), which therefore only expresses the Dfr-S isoform (C). Dfr-L is also expressed in adult fat body cells (H) and oenocytes (I). Scale bars represent 50 μm. (**J-L**) Immunoblot experiments using the Dfr-N antibody. Total protein was extracted from **(J)** BRGCs of female (F) and male (M) wandering L3 larvae or **(K)** BRGCs of synchronized L3 larvae, of indicated times (hours) after larval hatching (ALH). The concentration of Dfr-L decreased from early L3 to late L3. Actin was used as loading control. The Dfr-L and Dfr-S bands in BRGCs migrated more slowly than expected from the predicted molecular weight, most likely due to post-translational modifications. These bands were also present in blots from adult tissues, but there the bands with expected migration profile were dominating (Figure 1 –Figure supplement 1). (**L**) Quantification of relative protein concentration in panel (K).

### Larval to pupal transition is delayed in mutants that cannot produce Dfr-L

To study the *in vivo* function of Dfr-L, we generated *dfr* mutations using CRISPR/Cas 9-mediated genome editing. We isolated three different mutants carrying 1, 13 and 14 bp deletions downstream of the first in frame stop codon, and designated them *dfr^1^*, *dfr^13^*, and *dfr^14^* respectively (Figure 3A). Homozygous larvae of all three mutants displayed developmental delays, requiring between 5.5-7.5 days before pupariation, compared to five days for control (Figure 3B). Consequently, *dfr^1^*, *dfr^13^*, and *dfr^14^* adults were bigger than controls and their measured weight was increased (Figure 3C and 3D). We focused further work on the *dfr^14^* mutant, in which the 14 bp deletion removed part of a predicted RNA hairpin structure just 3’ of the stop codon, as well as causing a frameshift followed by numerous stop codons in the extending reading frame (Figure 3A). Immunostaining of ring glands with anti-Dfr-C did not produce any detectable staining in *dfr*^14^ (Figure 3E), as well as in other tissues conferring anti-Dfr-C immunostaining in control larvae and adults (Figure 2-figure supplement 1I), demonstrating that Dfr-L synthesis was abolished by the deletion. Conversely, Dfr-S translation was unaffected as both control and mutant stained positive for anti-Dfr-N (Figure 3E). This further suggests that the mutation neither affects *dfr* transcription negatively, nor translation until the first stop codon is reached. The abolished Dfr-L expression in *dfr^14^* BRGCs was confirmed on immunoblots (Figure 3F). As expected, loss of SCR also ensued a higher relative concentration of Dfr-S (Figure 3F), or possibly a severely truncated Dfr-L that likely would act as Dfr-S, as the isoforms are encoded from the same transcript. Taken together, these findings show that *dfr* SCR is necessary for correct timing of developmental transitions.

**Figure 3.**
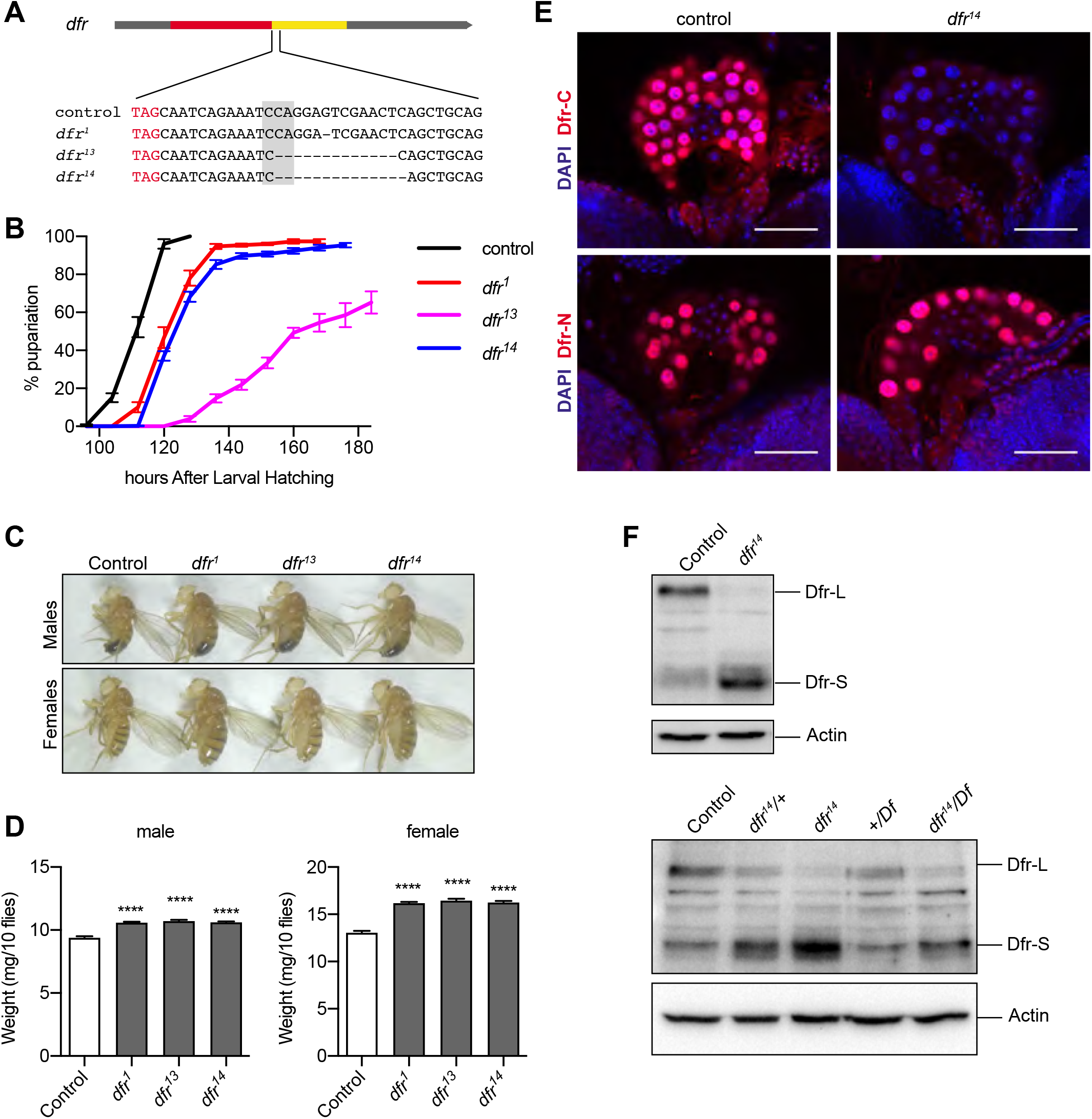
A 14-nt deletion downstream of *dfr* first in frame stop codon leads to developmental defects. (**A**) Upper panel, *dfr* gene structure; lower panel, nucleotide sequences of control *w^1118^* and *dfr* mutants. Three different *dfr* mutant strains were isolated, with 1, 13 and 14 nucleotide deletions downstream of the annotated stop codon, and named accordingly. The canonical stop codon is shown in red. The Protospacer Adjacent Motif (PAM) sequence targeted for cleavage by the CRISPR/Cas9 system is highlighted in grey. (**B**) The graph shows the percentage of pupariation relative to time in hours after larval hatching. The *dfr^1^*, *dfr^13^*, and *dfr^14^* mutations delayed the timing of pupariation compared to the *w^1118^* control. (**C**) Representative image of male and female adult flies of control *w^1118^*, *dfr^1^*, *dfr^13^*, and *dfr^14^*. (**D**) Quantification of fly weight of males and females. The *dfr^1^*, *dfr^13^*, and *dfr^14^* mutants show significantly increased body scale than controls when raised in noncrowded condition (30 larvae/vial). Data represent mean with SEM. n = 6 for all genotypes (males and females). Statistical analysis was performed using a One-Way ANOVA with Dunnett correction. ****, p<0.0001. (**E**) Immunostaining of control and *dfr^14^* brain/ring gland complexes with anti-Dfr-C (upper panels) and anti-Dfr-N (lower panels) antibodies. Dfr-L protein was not observed in *dfr^14^* prothoracic glands. Scale bars 100 μM. (**F**) Immunoblot of BRGC extracts incubated with anti-Dfr-N. Actin was used as a loading control. Upper panel: control and *dfr^14^*. The band corresponding to Dfr-L is barely detectable, while Dfr-S is considerably increased in *dfr^14^* mutants. Lower panel: *Df* is a large deficiency, *Df(3L)Exel6109*, which uncovers the *dfr* locus. Compared to the control, the Dfr-L band intensity is reduced in the *dfr^14^* mutant and *Df(3L)Exel6109* combinations, according to expectations, while the Dfr-S band shows increased intensity in *dfr^14^* background. Both Dfr-L and Dfr-S bands are weaker in +/*Df* compared to control extracts.

### The transcriptome is extensively dysregulated in larvae lacking the Dfr-L isoform

We reasoned that the C-terminal extension may provide Dfr-L with unique features in transcriptional regulation. To investigate this, RNA-seq analysis was applied to compare the transcriptome profiles in BRGCs (where SCR is very prominent) and in body tissues, separately, from *dfr^14^* 3^rd^ instar wandering larvae to those of controls. A two-dimensional scaling analysis based on the leading fold changes indicated a genotype-specific separation in the first dimension in both tissues (Figure 4A-B), supporting a specific role of Dfr-L in target gene regulation. The total number of differentially expressed targets (FDR<0.05) with a designated FBgn number (from hereon referred to as differentially expressed genes [DEGs]) was clearly larger in the BRGC than in the body (Figure 4C-D, Table S1), correlating well with the high rate of SCR in this tissues. In both groups, a slight majority of DEGs displayed increased expression. Several DEGs were strongly affected in the mutant, e.g. 53 in the BRGC and 82 in body had a log2±fold change >5 (Figure 4E-F). Gene ontology (GO) enrichment analysis of terms associated with biological processes revealed that the bulk of significant terms were linked to DEGs with reduced expression (down) in the BRGC of *dfr^14^* (Figure 4G), indicating a role for SCR and of the Dfr-L isoform in expression of these genes. These enriched terms encompassed diverse processes such as “positive regulation of gene expression”, “DNA replication initiation”, “protein deacetylation”, “sensory organ development”, “chromatin organization” and “Notch-signaling”, to name a few (Figure 4G; see Table S2 for the full list). For DEGs with increased expression in the BRGC in the *dfr^14^* mutant, enrichment and diversity were lower, but revealed some diverse processes (Figure 4G). In the body, only a few processes, associated with immunity and odor sensing, were significantly downregulated (Figure 4G). Enrichment analysis was also performed on terms related to molecular function (Table S2). In the BRGC and associated with DEGs with decreased expression, all enriched terms were related to DNA binding functions, including “transcription factor/cofactor activity” and “chromatin binding”. This implies that Dfr-L has broad downstream effects on the transcriptome, acting upstream of many other transcriptional regulators. The overall conclusion from the RNA-seq results is that SCR of *dfr* mRNA produces an alternative transcription factor isoform with uniques features and regulatory activity of a large number of target genes.

**Figure 4.**
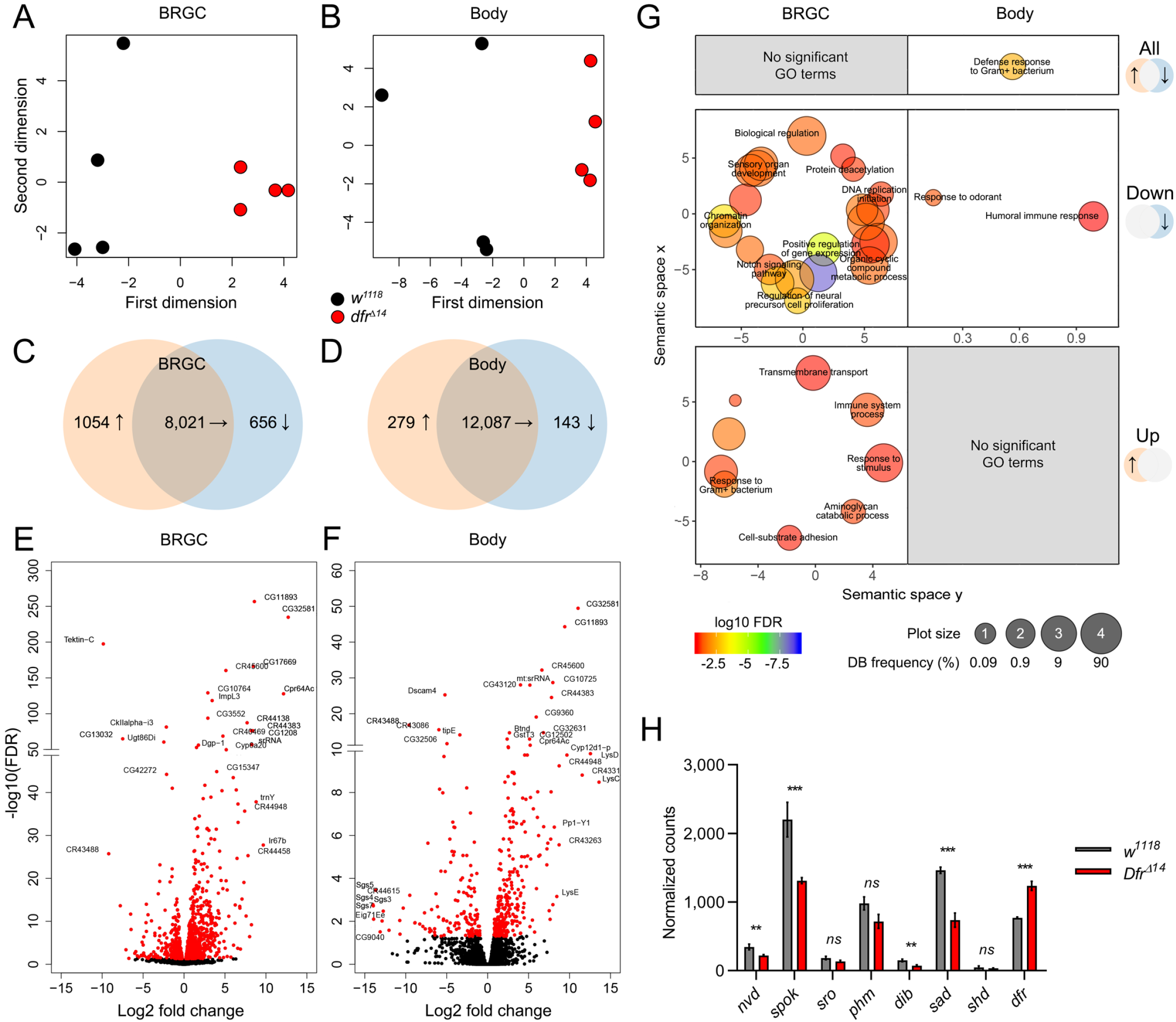
Absence of Dfr-L causes extensive transcriptional changes in the BRGC of wandering larvae. **(A-H)** RNA-seq analysis of BRGCs or bodies derived from control versus *dfr^14^*. Data were generated from all transcripts with a designated FBgn number and expressed in at least one of the groups in respective tissue. Each tissue was analysed separately. **(A-B)** Two-dimensional scatterplot depicting leading log2 expression differences between the transcriptomes of controls versus *dfr^14^* derived from either BRGC (A) or body (B). **(C-D)** Venn diagrams of unaltered, increased or decreased transcript levels in *dfr^14^* (FDR<0.05) in BRGC (C) and body (D). **(E-F)** Vulcano plots of all genes in BRGC (E) and body (F), expressed in at least two out the four groups. Differentially expressed transcripts are highlighted in red. **(G)** Gene ontology (GO) analysis was performed using GOrilla (terms with an FDR<0.1 were considered) and summarized in REVIGO to remove redundant terms (dispensability >0.5). Separate analyses were performed using either all (top panels), downregulated (middle panels), or upregulated (bottom panels) hits from respective tissue. Circle sizes represent GO term frequency (in log10-scale) in the underlying database, e.g. a small circle depicts a more specific term. Circle color and scale bar reflects log10 FDR-value. Scatterplot axes refer to semantic similarities between GO terms within a two-dimensional space (the values have no intrinsic meaning *per se*). For the complete list of GO terms see Table S2. For ease of viewing, the dispensability threshold was set to <0.2 for spelled out GO terms. **(H)** Comparative boxplot of BRGC expression of Ecdysone biosynthesis genes and *dfr*. Asterisks indicate differential expression between groups (FDR<0.01). Note that *shade (shd)* was included as a negative control since it is not normally expressed in the BRGC. For the complete list of ecdysone-associated genes, see Table S3.

It has earlier been shown using RNA interference (RNAi) that Dfr is involved in regulation of the ecdysone biosynthesis genes expressed in the prothoracic gland (PG, see below), *neverland (nvd), spookier (spok)*, *shroud (sro)*, *phantom (phm)*, *disembodied (dib)* and *shadow (sad)* (Cheng et al., 2014; Danielsen et al., 2014). In these studies, both *dfr* isoforms were targeted for knockdown as they are encoded from the same transcript. Thus, it was not possible from those RNAi-based experiments to deduce the discrete role(s) of Dfr-S and Dfr-L on expression of these target genes. Here, we found that expression of *nvd, spok, dib* and *sad* is reduced the BRGC of *dfr^14^* (Figure 4H). This indicates that the Dfr-L isoform is required for normal expression levels of these genes and, consequently, for steroidogenesis. Of note, expression levels of *dfr* mRNA *per se* was slightly, but significantly, increased (Figure 4H), possibly reflecting *dfr* autoregulation (Certel et al., 1996; Junell et al., 2010). We therefore ruled out that the observed effects on ecdysone biogenesis genes was caused by impaired *dfr* mRNA expression.

Defective ecdysone levels affect the temporal expression of ecdysone-responsive genes. Despite lacking the temporal aspect, the RNA-seq data revealed abolished expression of members of the salivary gland secretion family (*Sgs3*, *Sgs4*, *Sgs5, Sgs7, Sgs8*) and the ecdysone-inducible gene *Eig71Ee* in the body of *dfr^14^* (Figure 4F, Table 3). In summary, these findings show that the inability of *dfr^14^* mutants to produce the Dfr-L isoform by SCR has extensive effects on the transcriptome of wandering larvae. The transcriptome analysis further revealed a plausible requirement of *dfr* SCR for regulation of ecdysone biosynthesis and its downstream processes, which became the focus of our subsequent investigations.

### Expression of the steroidogenic enzymes Nvd, Dib and Sad is regulated by the readthrough product Dfr-L

In *Drosophila*, the neuroendocrine organs corpora allatum (CA), PG, and corpora cardiaca (CC) are fused into a compound structure, the ring gland (RG), which is attached to the brain (Figure 5-figure supplement 1A). To visualize the threedimensional (3D) structure of this endocrine organ, we performed 3D re-constructions based on a confocal stack of a BRGC (Figure 5-figure supplement 1B and Video 1). The PG is composed of the large ring gland lateral cells; CA cells are smaller, medial in the RG. The PG is the site for the ecdysone biosynthetic pathway with expression of all the enzymes required for the biosynthesis from cholesterol to ecdysone (Figure 5A). The last step of the ecdysone biosynthesis pathway, the conversion of ecdysone to the bioactive 20-hydroxyecdysone, takes place in peripheral tissues and is catalyzed by the enzyme Shade (Shd)(Petryk et al., 2003), and has not been reported to be a target of Dfr regulation.

**Figure 5.**
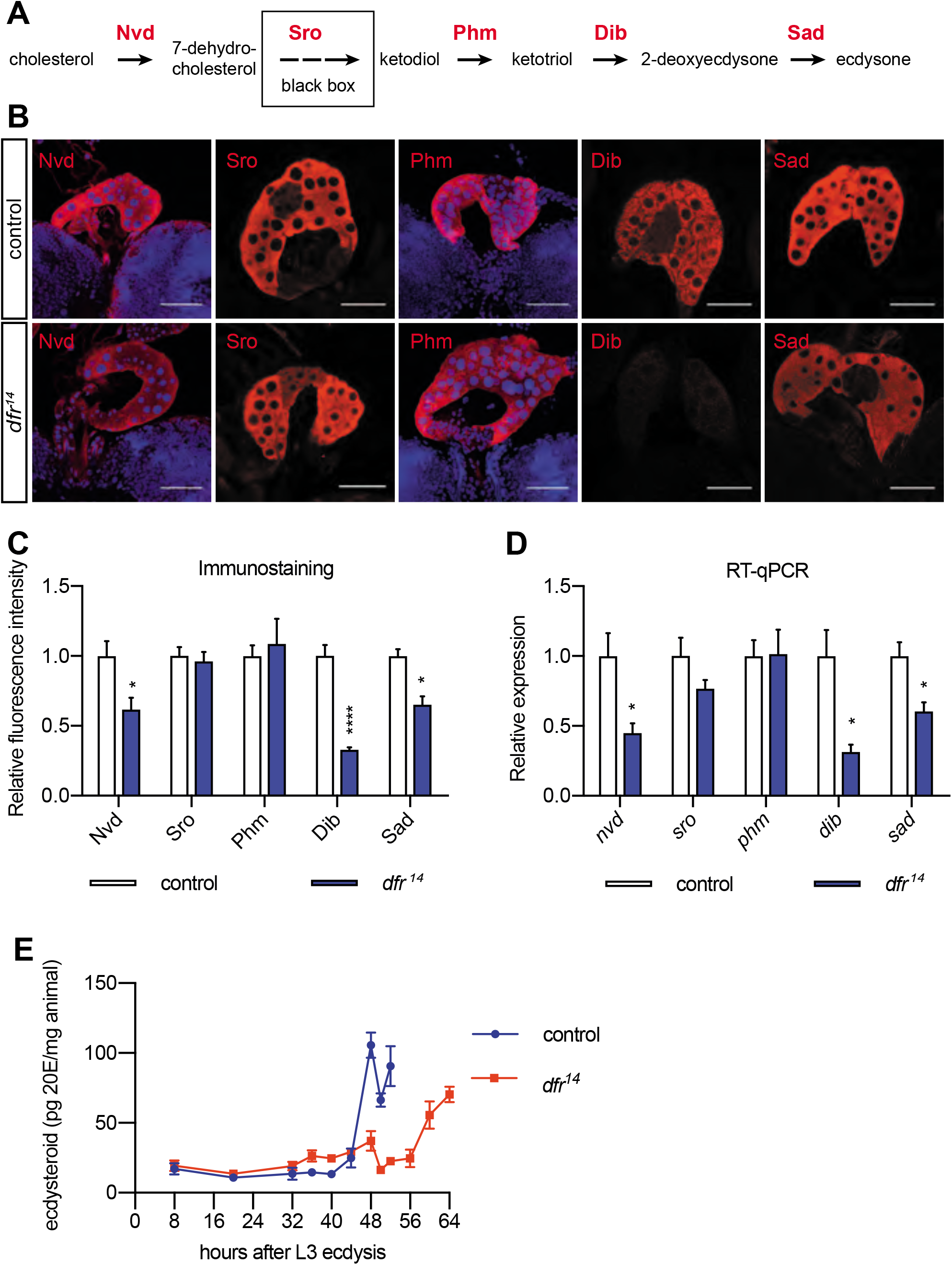
The expression of Nvd, Dib and Sad is compromised in the *dfr^14^* mutant. (**A**) A scheme showing ecdysone biosynthesis steps in the *Drosophila* prothoracic gland. Steroidogenic enzymes Nvd, Sro, Phm, Dib and Sad are sequentially required to convert cholesterol to ecdysone. (**B**) Immunostaining of control (upper panels) and *dfr^14^* (lower panels) BRGCs for the steroidogenic enzymes Nvd, Sro, Phm, Dib and Sad. The immunostaining of the enzymes is shown in red, DAPI staining in blue. Scale bars 50 μm, except for the images stained for Sad (100 μm in these two images). (**C**) Quantification of relative fluorescence intensity in (B). The expression of Nvd, Dib and Sad was significantly reduced in *dfr^14^* compared to control. (**D**) RT-qPCR results of the steroidogenic genes. Transcript levels of *nvd*, *dib* and *sad* were significantly decreased in *dfr^14^*. (**E**) Ecdysone biosynthesis is compromised in *dfr^14^* larvae. Ecdysone concentrations were quantified in a 2, 4 or 12-hour time window from early to late L3. *n* = 4 for each time point. In *w^1118^* control larvae, ecdysone peaks at around 48 h after L3 ecdysis during L3 development. In *dfr^14^* larvae, the ecdysone peak is delayed.

In line with the transcriptome data, immunostaining of *dfr^14^* mutant BRGCs showed that the expression of the steroidogenic enzymes Nvd, Dib, and Sad, but not Phm and Shd, was significantly decreased compared with control (Figure 5B-C). This was also confirmed by quantitative reverse transcriptase-PCR (qRT-PCR) (Figure 5D). To explore the functional importance of Dfr-L for temporal ecdysone production, we performed kinetic profiling of 20E titers in *dfr^14^* and control larvae after L3 ecdysis (AL3). As expected, control larvae showed a peak of 20E prior to pupariation, around 48h AL3. In *dfr^14^* mutant larvae, however, 20E titers remained low until around 60 h AL3 (Figure 5E). This demonstrates that SCR and the C-terminally extended Dfr-L isoform is required to ensure appropriate and dynamic ecdysone synthesis and release from the PG to properly time pupariation and metamorphosis.

### Overexpression of either Dfr-S or Dfr-L in the prothoracic gland causes developmental arrest

Next we analyzed the effects *in vivo* of targeted Dfr-S and Dfr-L overexpression, using independent *UAS-dfr-S* and *UAS-dfr-L* transgenic flies crossed with a temperature-sensitive Gal4 driver, *phm-Gal4^ts^* (*tub-Gal80^ts^; phm-Gal4*), for PG-specific expression of the two isoforms at specific times of development. The *dfr-S* construct carries sequences encoding ORF1 only (Figure 1A and 6A). The *dfr-L* construct was created by introducing a point mutation, converting the first TAG stop codon to AAG, thereby acting as an obligatory ORF1-ORF2 fusion transgene (Figure 6A). Overexpression of each isoform was confirmed in extracts from BRGCs dissected from synchronized late L3 larvae (Figure 6B). To our surprise, overexpression of *dfr-S* or *dfr-L* did not promote premature development, but instead phenocopied loss-of-function mutations in ecdysone biosynthesis genes, but at distinct developmental stages. Overexpression of *dfr-S* in the PG, led to developmental arrest at first larval instar (L1), a characteristic phenotype due to lack of ecdysone production, and also the phenotype of *dfr* RNAi (Danielsen et al., 2014). Partial rescue was observed upon 20E provision in the diet, as larvae developed into L2, but not L3 or pupae (Figure 6C-D), confirming that these larvae did not produce enough ecdysone. Overexpression of *dfr-L* in the PG also led to developmental arrest, but in L3 stage (Figure 6E-G), indicating that the ecdysone titers were appropriate in L1-L2 larvae, but not for pupariation. These L3 larvae continued to feed for more than 5 days, thereby gaining weight and volume (Figure 6F-G), and stayed at juvenile stage for up to one month until death (Figure 6E). Since there was no difference in volume between control and *dfr-L* overexpression larvae at day 5 after larval hatching (ALH), we concluded that larval growth rate was not affected *per se*, and that the primary phenotype is the inability to pupariate. Further analysis confirmed that overexpression of *dfr-S,* as well as *dfr* RNAi, had a strong negative effect on *nvd, phm, dib* and *sad* mRNA levels, while *dfr-L* overexpression showed less dramatic effects (Figure 6-figure supplement 1A-D). Thus, Dfr-S and Dfr-L confer different regulatory effects on downstream target genes.

**Figure 6.**
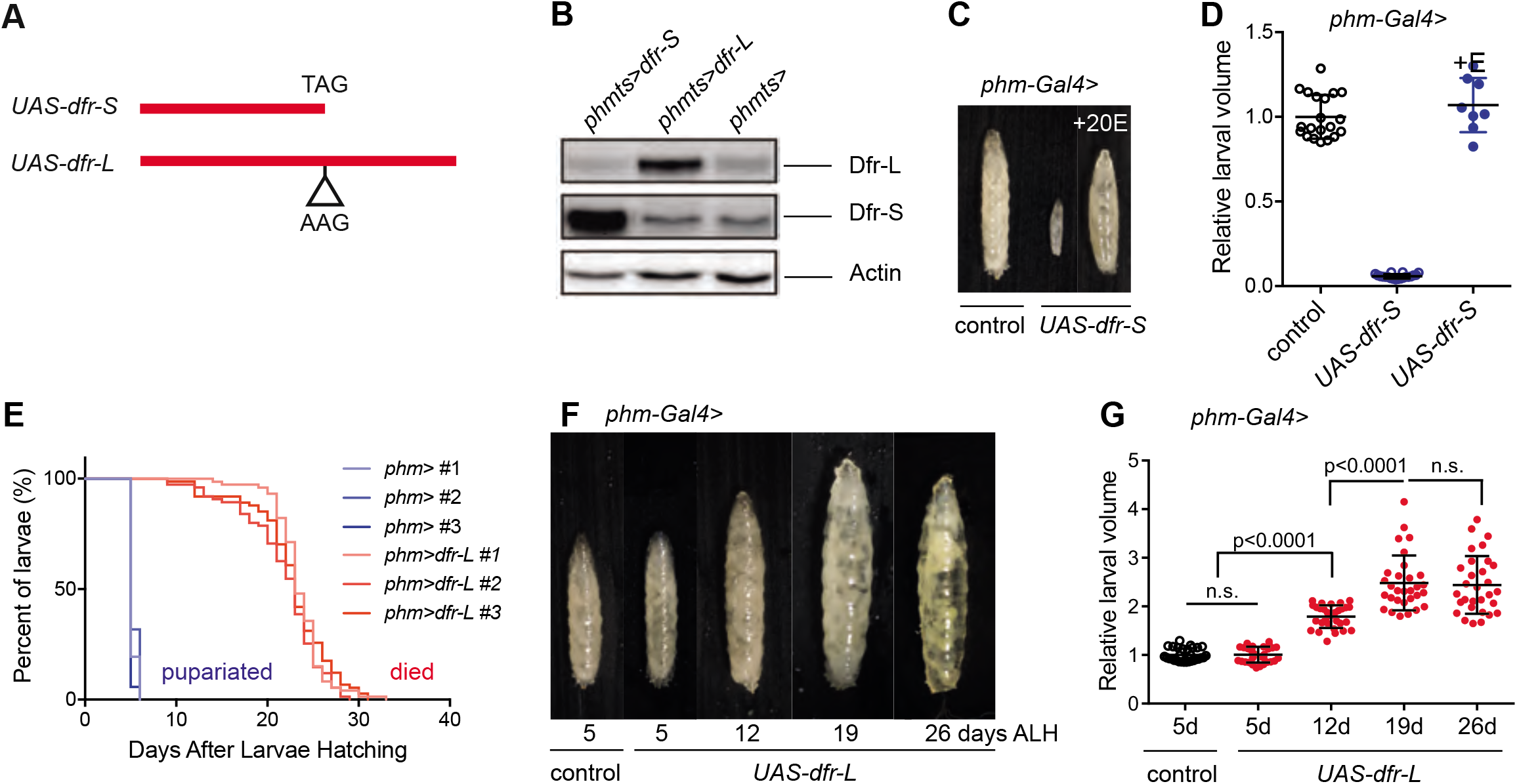
Dfr-S and Dfr-L overexpression causes arrest at different developmental stages. (**A**) Schematic drawing of expression constructs for Dfr-S and Dfr-L respectively. A point mutation was introduced to change the stop codon TAG to a lysine codon, AAG. (**B**) Overexpression of *UAS-dfr-S* and *UAS-dfr-L* in larval PG using a temperature-sensitive *Phm-Gal4* driver (*Phmts-Gal4).* Immunoblot using Dfr-N antibody. Proteins were extracted from three BRGCs for each genotype. Overexpressed Dfr forms migrated at the same position as endogenous ones. Actin was used as loading control. (**C-G**) Overexpression of *UAS-dfr-S* and *UAS-dfr-L* in larval PG using the *Phm-Gal4* driver. **(C-D)** Overexpression of Dfr-S in the PG led to arrest at first larval instar (L1). The developmental arrest was partially rescued by 20E feeding (+20E). Quantification of the relative larval volume (D). (**E**) Survival plot showing the percentage of larvae entering pupariation (control, blue) or eventually succumbing to death (Dfr-L overexpressing larvae). Control larvae (*phm-Gal4>*) pupariated at around 5-6 days after larval hatching. *phm-Gal4>UAS-dfr-L* led to L3 arrest. The larvae were not able to pupariate and died in the end. (**F**) Representative images of larvae at different days after larval hatching (ALH). *phm-Gal4>UAS-dfr-L* expression resulted in giant larvae. (**G**) Quantification of the relative larval volume in panel (E). There was no significant difference in larval volume between *phm-Gal4>UAS-dfr-L* and control larvae at 5 days ALH. The *phm-Gal4>UAS-dfr-L* larval volume increased with time.

### Clonal overexpression of Dfr-S depletes Dfr-L, leading to loss of ecdysone biosynthesis gene expression in a cell autonomous manner

To decipher the isoform-specific regulatory effects on steroidogenesis enzyme expression *in vivo*, we performed mosaic clonal analysis. We first analyzed Dfr immunostaining in GFP-marked *flp-*out clones overexpressing *dfr-S* or *dfr-L*. The nuclear anti-Dfr-N fluorescence intensity was significantly increased, while it was lost in *dfr-RNAi* clones, as expected (Figure 7A-B). Staining of *dfr-RNAi* and *dfr-L* clones with anti-Dfr-C showed a similar pattern (Figure 7C-D). Surprisingly, in *dfr-S* overexpressing clones, the anti-Dfr-C signal was gone, indicating that *dfr-S* overexpression led to depletion of Dfr-L in the PG (Figure 7C-D). This could either be due to suppression of endogenous *dfr* expression or by degradation of the Dfr-L protein. Using the same strategy, immunostainings of Nvd, Phm, Dib and Sad showed that reducing *dfr* expression by RNAi led to a significant reduction of all four proteins in GFP-labelled *flp*-out clones (Figure 7E-F and Figure 7-figure supplement 1A-F), but not in control clones, confirming the critical role of *dfr* in activation of *nvd, phm, dib* and *sad* genes (Danielsen et al., 2014). Overexpression of either *dfr-S* or *dfr-L*, also suppressed Nvd, Phm, Dib and Sad proteins expression in the PG clones (Figure 7E-F and Figure 7-figure supplement 1A-F), corroborating the conclusion that overexpression of *dfr-S* or *dfr-L* leads to ecdysone defects. Taken together, the marked decrease of several of the ecdysone biosynthesis genes after overexpression of *dfr-S* and *dfr-L* provides a likely explanation to the developmental arrest phenotypes presented in Figure 6. It further emphasizes the critical role of temporally controlled and balanced expression of Dfr-S and Dfr-L isoforms for normal development and progression through the ecdysone-regulated larval and pupal transitions.

**Figure 7.**
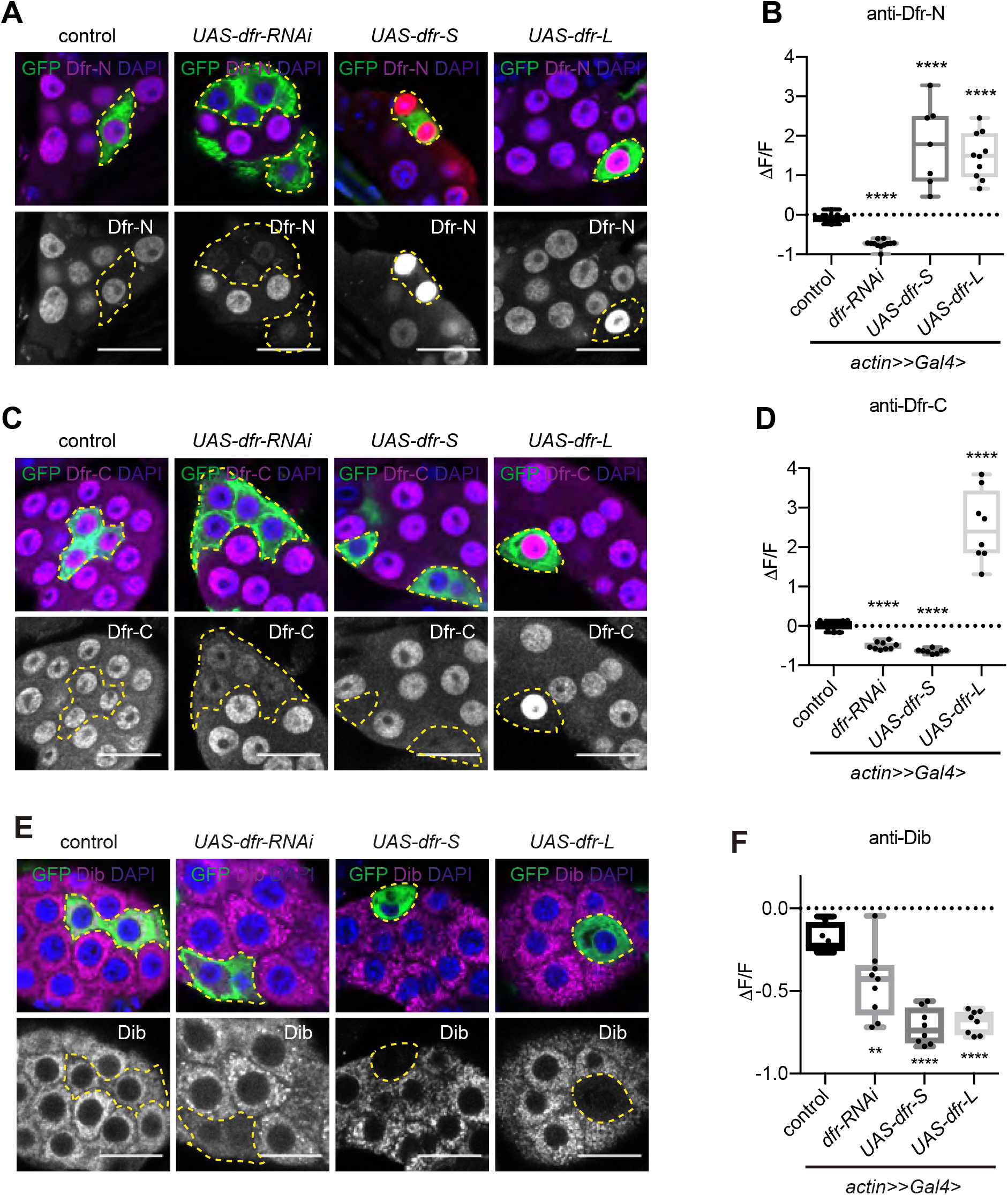
Dfr regulates the expression of the ecdysone biosynthesis gene *dib*. **(A)-(F)** Prothoracic glands (PGs) carrying GFP-labelled flp-out clones that express different transgenes (control, *UAS-dfr-RNAi*, *UAS-dfr-S*, and *UAS-dfr-L*). By applying a heat-shock at L1 stage, Flp recombinase activity was induced, which stochastically removed a stop cassette downstream of the *Actin* promoter in the tissue. Thereby *Actin-Gal4* expression was activated, which directed the expression of the target transgenes within the clone, including marking the clones with GFP. The PGs were stained with anti Dfr-N (A), Dfr-C (C) or Dib (E) antibody, and the clones are outlined with yellow dashed lines. **(A)** Upper panels: anti-Dfr-N staining is shown in red, GFP in green, DAPI stains nuclei in blue. Lower panels: Dfr-N staining shown in grey. GFP expression did not affect Dfr protein expression in the control. Downregulation of *dfr* mRNA by *UAS-dfr-RNAi* reduced Dfr protein level in the PG clones. Both *UAS-dfr-S* and *UAS-dfr-L* expression increased Dfr immunostaining in the clones. (**B**) Quantification of relative fluorescence changes in the clones in panel (A). ΔF/ F measured the change of fluorescence intensity between a clone cell and a neighbor cell. (**C**) Upper panels: anti-Dfr-C staining is shown in red, GFP in green, DAPI stains nuclei in blue. Lower panels: anti-Dfr-C staining is shown in grey. GFP expression did not affect Dfr-L protein expression in the control. Downregulation of *dfr* mRNA decreased Dfr-L protein levels in the PG clones. Overexpression of the short form Dfr-S decreased Dfr-L protein levels in the clones, while *UAS-dfr-L* increased Dfr-L abundance. (**D**) Quantification of relative fluorescence changes in the clones in panel (C). (**E**). Upper panels: Anti-Dib staining is shown in magenta, GFP in green, DAPI stains nuclei in blue. Lower panels: anti-Dib staining is shown in grey. Dib expression decreased in *UAS-dfr-RNAi* clones. Dib protein expression was also diminished in *UAS-dfr-S*, and *UAS-dfr-L* overexpression clones. Scale bars, 25 μM. (**F**) Quantification of the relative fluorescent changes in panel (E).

### Dfr-L and Molting defective (Mld) synergistically activate transcription of ecdysone biosynthesis genes

To determine whether the C-terminal extension of Dfr-L promotes transcriptional activation or repression we performed luciferase reporter assays in *Drosophila* S2 cell cultures. We focused on two of the ecdysone biosynthesis genes, *nvd* and *spok*, whose expression was hampered in *dfr^14^* BRGCs (Figure 4N). These genes also contain putative binding sites for Dfr (Danielsen et al., 2014). Surprisingly, transfection with *dfr-S*, *dfr-L* or both, repressed *nvd-Fluc* (Figure 8A) although Dfr-L is required for its expression (Figure 4N), and *spok-Fluc* was not significantly affected (Figure 8B). This is, however, in line with the *in vivo* results showing that overexpression of Dfr-S and Dfr-L can inhibit ecdysone biosynthesis. It has previously been reported that *nvd* and *spok* are regulated by the zinc-finger transcription factor Molting defective (Mld) (Danielsen et al., 2014; Neubueser et al., 2005; Uryu et al., 2018). In accordance with those studies, expression of Mld activated *nvd-Fluc* and *spok-Fluc* approximately 5fold (Figure 8A-B). Strikingly, cotransfection with *dfr-L* and *mld* expression plasmids, both *nvd-Fluc* and *spok-Fluc* reporters were synergistically activated (approximately 10-fold and 50-fold respectively) (Figure 8A-B), indicating a coordinated role of Dfr-L and Mld in regulation of *nvd* and *spok* expression. Converesely, cotransfection with *dfr-S* and *mld* instead hampered *nvd-Fluc* expression compared to *mld* transfection alone, whereas *spok-Fluc* was induced but to a less degree than *dfr-L* and *mld* (Figure 8A-B). In conclusion, the C-terminal extension of Dfr by SCR enhances its *trans-*activation capacity in conjunction with Mld.

**Figure 8.**
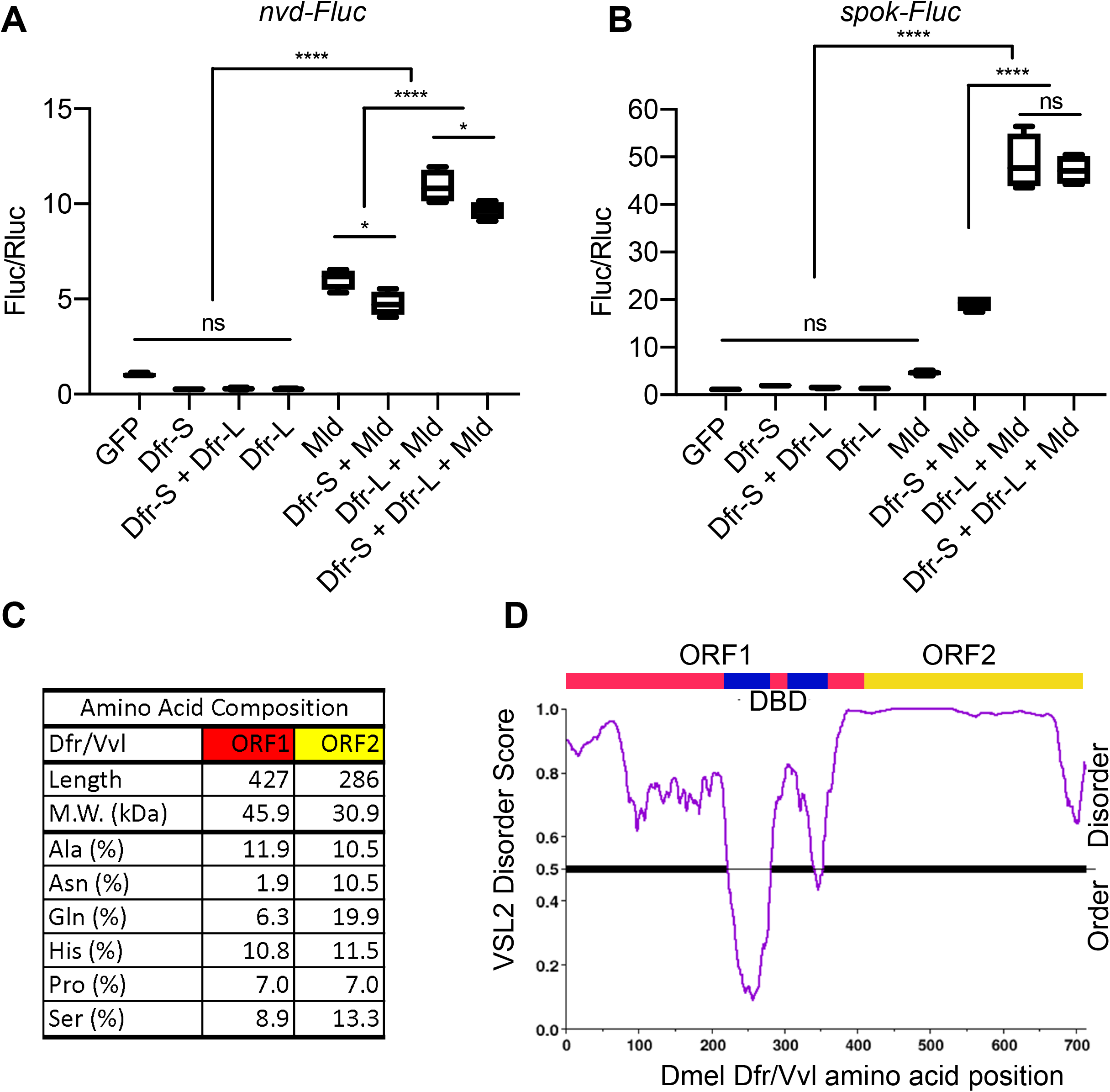
The SCR-dependent extension of Dfr-L acts as a transcriptional activation domain and is highly disordered. **(A-B)** Transcriptional reporter assays with *nvd-Fluc* (A) and *spok-Fluc* (B) expression in response to expression of Dfr-S, Dfr-L and Mld independently, and in combination, in *Drosophila* S2 cells. GFP expression plasmid was used as a negative control. Y axis shows the luminescence of firefly luciferase over renilla luciferase (Fluc/Rluc). *N*=4. *p<0.05; ***p<0.001; ****p<0.0001. **(C)** Amino acid composition of Dfr ORF1 and ORF2, showing the six most prevalent amino acids in ORF2. Especially Asn, Gln and Ser are more frequent in ORF2 than in ORF1. **(D)** Disorder analysis of *Drosophila melanogaster* Dfr-L. The intrinsic disorder of Dfr-L was calculated by the VSL2 algorithm (http://www.pondr.com/). Schematic representation of ORF1 (red), ORF2 (Yellow) and the DNA-binding domains (DBD, blue) are shown above the disorder graph. The horizontal bold line indicates 0.5 disordered score, above which the amino acid sequence is disordered.

**Figure 9.**
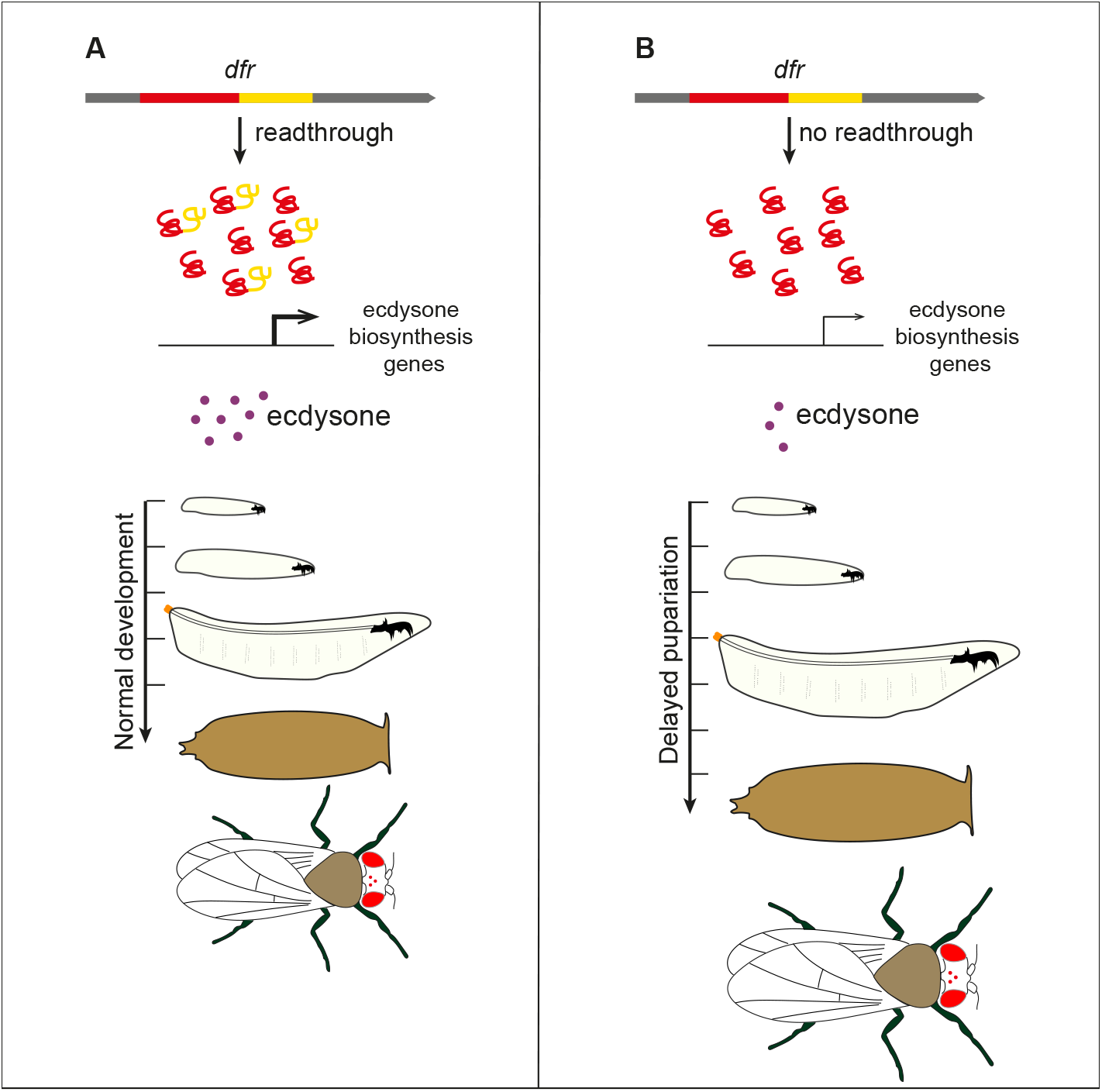
Translational stop codon readthrough of *dfr* regulates ecdysone biosynthesis and *Drosophila* development. Dfr is highly expressed in the *Drosophila* PG. Conventional translation of *dfr* generates the short form Dfr-S, while SCR produces Dfr-L. Both Dfr-S and Dfr-L regulate ecdysone biosynthesis genes and are involved in *Drosophila* development. High levels of *dfr* SCR in the PG is required for the proper production of ecdysone and developmental transitions. Without *dfr* SCR (e.g., in *dfr^14^*), ecdysone biosynthesis is compromised, which leads to prolonged larval development, delayed pupariation and increased adult size.

### The C-terminal extension of Dfr consitutes an evolutionarily conserved intrinsically disordered region

Having established that Dfr-L can be a stronger transcriptional activator than Dfr-S, we sought for an underlying explanation to this difference in the amino acid sequence and properties of the SCR-derived C-terminal extension of Dfr-L. Searches for putative functional domains by InterPro and ELM within the C-terminal extension did not provide any high fidelity hits. The C-terminal extension has, however, an unusually high proportion of the amino acids Gln (19.9%), Ser (13.3%), His (11.5%), Ala (10.5%), Asn (10.5%) and Pro (7.0%) (Figure 8C). It also contains several low complexity regions with long stretches of Gln, Asn and His/Pro (Figure 1C). Such low complexity regions are frequently observed in *trans-*activation domains (tADs) of eukaryotic transcription factors (Arnold et al., 2018; Mitchell and Tjian, 1989), and it also suggests the presence of intrinsically disordered regions (IDRs). This was confirmed using IUPred and PONDR analytical tools, which predicted that the entire C-terminal extension is disordered (Figure 8D). Interestingly, similar amino acid composition (Figure 8, figure supplement 1A) and the presence of a large IDR was evident in the predicted SCR-derived C-terminal extensions of *dfr/vvl* in a number of dipteran species (Figure 8, figure supplement 1B-H). Thus, the physico-chemical properties of the C-terminal extension seems to be more important and under positive natural selection, than the presence of specific protein domains or sequence identity.

## Discussion

Programmed, alternative decoding of the genome, such as SCR and translational framshifting have recently got increased attention through comparative genomics analyses and ribosome profiling experiments, indicating that alternative coding is pervasive and evolutionarily conserved (Dunn et al., 2013; Jungreis et al., 2016; Jungreis et al., 2011; Loughran et al., 2018; Rodnina et al., 2019; Sapkota et al., 2019). In the present study, we provide several lines of evidence to show that *dfr* mRNA undergoes SCR in *Drosophila*. Firstly, the pull-down of a Dfr-L-Myc fusion protein confirmed that Myc was properly translated as a result of SRC of *dfr* mRNA. Secondly, the mass spectrometry identified peptides that matched the C-terminal extension sequence of Dfr. Thirdly, we show that the stop codon UAG was decoded as glutamine. Lastly, immunostaining in larvae and adults with an antibody that recognizes the Dfr C-terminal extension confirmed SCR of *dfr* mRNA *in vivo*. Strikingly, the readthrough rate of *dfr* is as high as 50% in certain tissues during specific stages of development, indicating that *dfr* SCR is a regulated event with functional consequences. We also show that Dfr-L plays an important developmental role in modulating steroidogenesis, thereby affecting the temporal progression through the larval developmental stages to pupariation and metamorphosis.

A few studies using ribosomal protection assays in insects and human cells have shown that the rate of SCR differs between tissues and cell types, suggesting that SCR is a programmed and regulated process (Dunn et al., 2013; Pancsa et al., 2016; Sapkota et al., 2019). However, a clear link between the ribosomal profiling data and a functional importance *in vivo* of SCR has essentially been lacking in metazoans, and it has been argued that SCR is generally nonadaptive (Li and Zhang, 2019). Our work shows that *dfr* SCR differed in a stage- and tissue-specific manner (Figure 2), strongly indicating the involvement of *trans*-acting factors, such as protein, RNA or other molecules, interacting with *cis-*acting elements in the affected genes. To identify such *cis*- and *trans-*acting molecules and to elucidate the underlying mechanisms of this regulation will be an important undertaking in future work.

A deletion that abolished the C-terminal extension had strong effects on the transcriptional profile of *dfr^14^* mutant larvae, with pronounced effects on genes involved in gene expression, neural proliferation, sensory organ development, and immune system processes. This together with the hampered ecdysone production and developmental delays of *dfr^14^* mutant larvae demonstrates the importance of SCR and the C-terminally extended Dfr-L isoform *in vivo*. Importantly, Dfr-L, but not Dfr-S, activated *nvd-Fluc* and *spok-Fluc* reporters in a synergictic manner together with Mld. Thus, the SCR-derived C-terminal extension and the Dfr-L isoform modulate the expression of steroidogenic enzymes, and in its absence, the developmental transitions between different life-cycle stages are delayed.

The first step in ecdysone production is the conversion of dietary cholesterol to 7-dehydrocholesterol (7DC), regulated by Nvd. Remarkably, *nvd* and *spok* are located in the pericentromeric regions thought to form constitutive heterochromatin (Fitzpatrick et al., 2005; Ono et al., 2006; Yoshiyama et al., 2006). Expression of heterochromatic genes has been suggested to require epigenetic regulators that control heterochromatic silencing, for example HP1a, and other chromatin remodeling complexes (Marsano et al., 2019). In this context, it is interesting to note that the transcriptome analysis of the *dfr^14^* mutant lacking Dfr-L, revealed that expression of genes involved in biological processes defined as “chromatin organization”, “protein deacetylation” and “positive regulation of gene expression” were reduced in *dfr^14^* mutant BRGCs. This suggests that Dfr-L may target genes, such as *nvd* and *spok*, at several levels, both directly in transcriptional activation, as well as indirectly by controlling epigenetic and chromatin modifying factors, and other transcriptional regulators. It is tempting to speculate that the Dfr-L isoform could play a distinct role in regulation of the heterochromatic genes *nvd* and *spok* by interacting with heterochromatin regulators.

The C-terminal extensions produced by SCR have in some cases been found to possess functional modules, e.g., peroxisomal localization signals (Freitag et al., 2012) and nuclear localization signals (Dunn et al., 2013). Both Dfr-S and Dfr-L are constitutively nuclear (Figure 2), indicating that SCR of *dfr* does not affect its subcellular localization. Both isoforms carry the same DNA binding POU and Homeo domains (Figure 8D), and both can act as transcriptional activators in reporter assays (Figure 8A and B). However, Dfr-L conferred stronger transcriptional activation of the luciferase reporters than Dfr-S, suggesting that the C-terminal extension of Dfr-L contains additional *trans-*activation domain(s) (tADs), and/or protein-protein interaction domains that favour interactions with transcriptional co-activators or chromatin modifiers.

Interestingly, the extension contains several low complexity regions, together constituting an intrinsically disordered region (IDR) (Figure 8C and D). Regions enriched for individual amino acids including glutamine, asparagine, histidine, serine, proline, alanine are well known to be abundant in different classes of tADs (Mitchell and Tjian, 1989). Similar amino acid composition was also evident in the predicted C-terminal extensions of Dipteran Dfr/Vvl proteins (Figure 8-figure supplement 1A-H), suggesting that the physico-chemical properties of Dfr C-terminal extensions are under purifying selection, and more important than the primary amino acid sequence *per se.* The presence of glutamine-rich regions is especially intriguing since this feature has repeatedly been linked to tADs (Gemayel et al., 2015; Gerber et al., 1994).

Low complexity regions and IDRs have recently been linked to liquid-liquid phase separation of transcription regulatory complexes (Boija et al., 2018; Chong et al., 2018). In a computational analysis of *Drosophila melanogaster* SCR candidate proteins it was found that the C-terminal extensions were significantly enriched in disordered and low complexity regions (Pancsa et al., 2016) raising the possibility that these in fact constitute regulatory entities that are added as C-terminal extensions through SCR. For transcriptional regulators like Dfr, for which the SCR is regulated in a spatiotemporal manner, the addition of an IDR/tAD to its C-terminus is therefore likely to have a major impact on a number of processes. The transcriptome profile of the *dfr^14^* mutant (Figure 4) support that hypothesis and indicates that SCR of *dfr* is important for neurogenesis, sensory organ development, the immune response and metabolic processes. Similarly, SCR of the mammalian gene for Argonaute1 (Ago1) produces the Ago1x isoform, which acts as a competitive inhibitor of the miRNA pathway, leading to increased global translation as a result of SCR (Singh et al., 2019). We conclude that SCR of regulatory proteins may play a more prominent role in controlling biological processes than previously anticipated.

## Material and Methods

### Fly stocks

Flies were maintained on potato medium (Dantoft et al., 2016) at 25 °C unless otherwise indicated with a 12 h light 12 h dark cycle. The *w^1118^*, *dfr* deficiency line Df(3L)Exel6109 (BL7588), *Aug-Gal4* (BL30137), *tub-Gal80ts* (BL7019), *UAS-mCherry* (BL38425), *Sgs3-GFP* (BL5885) and *vasa::Cas9* (BL51323) were obtained from Bloomington Drosophila Stock Center (BDSC). The *UAS-dfr-RNAi* line was provided by Sarah Certel; *UAS-dfr-S* and *UAS-dfr-L* transgenic lines are described below; *phm-Gal4*, was provided by Kim Rewitz.

### Phylogenetic analysis

Multiple sequence alignments of Dfr from selected species were performed using MAFFT (Katoh and Standley, 2013). In cases were SCR was not annotated, the open reading frame immediately downstream of the first stop codon, and in frame, was manually translated into amino acid sequences until the subsequent stop codon to achieve a hypothetical protein extension. The output were used to construct Phylograms in Simple Phylogeny (Saitou and Nei, 1987) using default parameters including the Neighbour-joining method and visualized by real branch lengths. Alignments were additionally imported into MView (Brown et al., 1998) to obtain the degree of consensus per base.

### Analysis of putative splicing or editing events

RNA was isolated from male flies using TRIzol (Invitrogen) and treated with DNase (Applied Biosystems) according to the manufacturer’s instructions. The isolated RNA was used for cDNA synthesis using the Access RT-PCR system (Promega) with *AMV* reverse transcriptase, and with primers amplifying a 500 bp region surrounding the first stop codon. The PCR products were run on agarose gel electrophoresis and analyzed using BioRad UV-vis camera. Thereafter the agarose gel band was excised from the gel, DNA extracted using QIAquick gel extraction kit (Qiagen), and used as template for DNA sequencing (Eurofins MWG Operon sequencing service).

### Gateway cloning

Different *dfr* expression constructs were made using a 3.7 kb full length *vvl/dfr* cDNA (provided by W. Johnsson) as template and pENTR^TM^ directional TOPO^®^ cloning kit according to the manufacturer’s instruction (Invitrogen). The following constructs were made: *dfr-3* construct contains 1284 bp cDNA sequence from the start codon to the first TAG stop codon, and can solely express Dfr-S; *dfr-4* construct contains the 2142 bp cDNA sequence from the start codon to the second TAG stop codon. It still carries the first stop codon and can express both Dfr-S and Dfr-L, the latter as a result of readthrough. To create an obligate Dfr-L expressin construct (*dfr-5)*, a point mutation was inserted in *dfr-4*, by inverse PCR with phosphorylated primers, converting the first in frame TAG stop codon to a lysine codon AAG. The *dfr-6* construct contains the coding sequence between the first and second stop codons (nt 1953 –2810), enabling expression of the 285 amino acid C-terminal extension for antibody production.

The following primers were used (5’-3’): *dfr-3*, *dfr-4, dfr-5* forward: CACCATGGCCGCGACCTCG *dfr-6* forward: CACCCAATCAGAAATCCAGG *dfr-3* reverse: GGCCGCCAACTGATGCGCCG *dfr-4, dfr-5, dfr-6* reverse: TTCGCCACCCGCTCCGCCCG The following primers were used to induce the point mutation (5’-3’): Forward primer: ⓟ-AAGCAATCAGAAATCCAGGAG Reverse primer: ⓟ-GTGGGCCGCCAACTGATGCG Destination plasmids for expression of untagged and tagged constructs of each isoform in cell cultures, bacteria and for P-element mediated transformation were made via recombination using the Gateway^®^ LR Clonase Enzyme mix according to the manufacturer’s instruction (Invitrogen).

### P-element mediated transformation

P-element-mediated transformation was performed according to Rubin and Spradling (Rubin and Spradling, 1982). The *pUAS-Dfr-S* and *pUAS-Dfr-L* plasmids were injected together with the Δ2-3 helper plasmid into the recipient strain, *yw* (Laski et al., 1986). The eclosed G0 flies were back-crossed with the *yw* flies, and G1 flies were crossed with balancer lines individually to establish stable transformant strains.

### CRISPR /Cas9 gene editing of *dfr/vvl*

The gene editing of *dfr/vvl* was performed using single gRNA according to (Port et al., 2014). Genomic DNA was isolated from the recipient fly strain *vasa::Cas9* line (BL51323). A region of 563 bp around the first in frame stop codon of the *dfr* gene was amplified by PCR and sequenced to determine potential polymorphism between *vasa::Cas9* line and the reference genome. Microinjections were carried out with 500 ng/μl gRNA plasmid. Injected G0 males were crossed with *w;; MKRS/TM6b* balancer stock, 2-3 progeny males from each cross were crossed with *w*;; *MKRS/TM6B* virgins. Stocks were established from the progeny. Homozygous larvae from each stock were chosen for genotyping. Initial experiments and the RNA sequencing was done with homozygous *dfr^14^* that had been outcrossed to *w;; MKRS/TM6B.* To further clean up the third chromosome, the *dfr^14^* mutant was crossed with *w^1118^,* and F1 females were outcrossed to *w^1118^* background for six generation. Genotyping was performed to trace the mutation in *dfr.*

Oligos for analysis of polymorphism and genotyping (5’-3’) were: Forward: CAGAAGGAGAAGCGCATGAC Reverse: TGCTGCTGGTGGTGTTTAAC. Oligos for gRNA plasmid (5’-3’) Forward: GCTGCTGCAGCTGAGTTCGACTCC Reverse: GGAGTCGAACTCAGCTGCAGAAAC

### Immunoprecipitation, in-gel digestion and mass spectrometry analysis

*Drosophila* S2 cells were transfected with 3μg of *pAWM-dfr4* using Effectene transfection kit (Qiagen) according to the manufacturer’s instruction. Transfected cells were harvested on day 4 after transfection, washed 2 times in PBS, homogenized in lysis buffer containing 20 mM Tris pH 7.8, 150 mM NaCl, 10 mM MgCl_2_, 2 mM EDTA, 10% Glycerol, 0.5% NP40, 1 mM DTT and protease inhibitor cocktail according to the manufacturer’s instruction. The homogenate was shaken gently at 4°C for 10 minutes and then centrifuged at 1500 g. Immunoprecipitation was done using mouse anti-Myc antibody (4A6, Millipore) at 1-3 mg/ml and Dynabeads^®^ Protein G (Thermo Fisher Scientific) according to the manufacturer’s instruction.

Eluted proteins were separated by 7.5% SDS-polyacrylamide gel electorphoresis. The band corresponding to Dfr4-Myc was excised manually from a Coomassie-stained gel. In-gel digestion, peptide extraction, MS analysis and data-base searches for protein identification were carried out at the Proteomics Biomedicum, Karolinska Institute, Sweden, as follows: In-gel digestion of the gel pieces were done using a MassPREP robotic protein-handling system (Waters, Millford, MA, USA). Gel pieces were destained twice with 100μl 50mM ammonium bicarbonate containing 50% acetonitrile at 40°C for 10 min. The protein was reduced by 10mM DTT in 100mM Ambic for 30 min and alkylated with 55mM iodoacetamide in 100mM Ambic for 20 min followed by in-gel digestion with 0.3 mg chymotrypsin (modified, Promega, Madison, WI, USA) in 50 mM ammonium bicarbonate for 5 h at 40°C. The tryptic peptides were extracted with 1% formic acid/ 2% acetonitrile, followed by 50% acetonitrile twice. The liquid was evaporated to dryness and the peptides were injected onto the LC-MS/MS system (Ultimate^TM^ 3000 RSLCnano chromatography system and Q Exactive Plus Orbitrap mass spectrometer, Thermo Scientific). The peptides were separated on a homemade C18 column, 25 cm (Silica Tip 360μm OD, 75μm ID, New Objective, Woburn, MA, USA) with a 60 min gradient at a flow rate of 300nl/min. The gradient went from 5-26% of buffer B (2% acetonitrile, 0.1% formic acid) in 55 min and up to 95% of buffer B in 5 min. The effluent was electrosprayed into the mass spectrometer directly via the column. The spectra were analyzed using the Mascot search engine v. 2.4 (Matrix Science Ltd., UK).

### In vitro transcription/translation

In vitro transcription/translation of *vvl/dfr* vDNA was carried out using TNT coupled reticulo lysate system (Promega) and T3 RNA polymerase (New England Biolabs Inc.) with *dfr* full-length cDNA in pBSKS (gift from W. Johnsson) as template, according to the manufacturer’s instruction.

### Antibody production

Antibodies against Dfr-C/ORF2 were raised in rat against a purified recombinant Dfr ORF2 protein produced in *E.coli.* Recombinant protein expression, purification, and immunization of rats were carried out by Agrisera AB, Vännäs, Sweden, as follows: GST-tagged Dfr-ORF2 protein was produced in BL21(DE3) and purified by affinity chromatography on a Glutathione Sepharose 4B column. The GST part was cleaved off from the recombinant protein using PreScission Protease (GE Healthcare Life Sciences) according to the manufacturer’s instructions. Purified Dfr-ORF2 protein without the tag was used for immunization of rats. Serum titers were analyzed by immunoassays and antibody specificity against Dfr-L using immunoblot assays.

### Immunocytochemistry of *Drosophila* tissues

*Drosophila* larvae were dissected in phosphate-buffered saline (PBS, pH 7.0) and fixed in 4% paraformaldehyde for 30 min at room temperature. The specimens were washed in PBST (PBS with 0.3% Triton X-100) three times, then blocked in PBST with 0.5% normal goat serum for 1 hour at room temperature. Antibody dilutions used were as follows: rat anti-Dfr-N (1:400), rat anti-Dfr-C (1:400), guinea pig anti-Neverland (1:1,000), guinea pig anti-Shroud (1:1,000), rabbit anti-Phantom (1:400), rabbit anti-Disembodied (1:400), and rabbit anti-Shadow (1:400). Secondary antibodies were Alexa Fluor 594 conjugated goat anti-rat (1:500), goat anti-rabbit (1:500), and goat anti-guinea pig (1:500). DAPI was used to stain the nuclei. Flp-out clones were also analyzed using this protocol.

### Immunoblot assays

Protein extraction from dissected tissues was performed as previously described (Dantoft et al., 2013). Extracts were separated by electrophoresis in a 10% SDS-polyacrylamide gel at constant current of 120 volt. Proteins were transferred to polyvinylidinefluoride membranes (Millipore Corporation, Billerica, MA, USA), subsequently blocked 5% dry milk in TBST (Tris Buffered Saline with 0.1% Tween 20) for 1 h at room temperature and then incubated with anti-Dfr-N, anti-Phm, anti-Dib, or anti-Actin (mAbcam 8224) as primary antibodies, and with ECL^TM^ anti-rat IgG (Amersham), ECL^TM^ anti-mouse IgG (GE Healthcare), and ECL^TM^ anti-rabbit IgG (GE Healthcare) as 2^nd^ antibodies. The blot was developed using either SuperSignal^TM^ West Femto maximum sensitivity substrate or SuperSignal^TM^ West Pico PLUS Chemiluminescent Substrate (Thermo Scientific) according to the manufacturers’ instructions. Digital images were acquired with ChemiDoc™ Imaging Systems (Bio-rad). Protein levels were quantified with Image Lab™ Software (Biorad) and normalized agaist Actin or Lamin. Statistics was performed using two-way ANOVA.

### RNA sequence analysis

Total RNA was extracted from BRGCs and bodies of wandering L3 larvae. The body samples were devoid of BRGCs, mouth hooks and salivary glands. The BRGC and body samples were collected from different larvae respectively and hence considered as separate experiments. Four biological replicates were prepared for each group. The RNA samples were further cleaned up with Qiagen RNeasy kit (Qiagen, Valencia, CA) according to the manufacturer’s instructions. Sequencing was performed at Science for Life Laboratory (National Genomics Infrastructure, Stockholm node), using a HiSeq2500 (Illumina TruSeq Stranded mRNA) with Poly-A selection. Raw data in binary base call (BCL) format were converted to FastQ using bcl2fastq_v2.19.1.403 from the CASAVA software suite. All samples passed the quality test pipeline. Mapped reads per gene (ENSEMBL BDGP6 assembly) were quanitified using featureCounts. Datasets from body and BRGC were analysed separately. Genes with no counts in either group of respective tissue were filtered out from the analysis (7,827 in BRGC; 5,045 in body) resulting in 9,731 (BRGC) and 12,513 (body) remaining. Differences in library sizes between samples where accounted for using the calcNormFactors function to scale reads according to the effective size of each library. Annotation was performed using the Bioconductor 3.8 annotation package org.Dm.eg.db. Differential expression analysis was carried out using Bioconductor 3.8 with the edgeR 3.8 package in R 3.5.2 according to the edgeR user’s guide (26 October 2018 revision). Exact test was applied with a false discovery rate (FDR) threshold set to <0.05 for significant hits. Multidimensional scaling (MDS) of samples was plotted using edgeR using the default setting of leading logfold-changes between each pair of sample to map the corresponding distances. Venn diagrams were constructed in Photoshop CC 2015. Vulcano plots were constructed using the ggplot2 package in R. Gene ontology (GO) analyses were performed in BRGC or body, respectively, using GOrilla (FDR<0.1 was considered significant) (Eden et al., 2007; Eden et al., 2009). As background gene list, all enlisted IDs with expression in at least one of the groups in respective tissue was used. Analyses were performed on upregulated, downregulated or all differentially expressed hits separately. Redundant GO terms were filtered out using REVIGO (Supek et al., 2011) with allowed similarity set to “low” (dispensability <0.5). Generated REVIGO scripts for semantic scatterplots were imported to RStudio for plotting.

### Quantitative RT-PCR

Female virgins of *tub-Gal80^ts^; phm-Gal4* (200-300 virgins in each bottle) were crossed with *w^1118^, UAS-dfr-RNAi, UAS-dfr-S*, and *UAS-dfr-L*, respectively. Embryos were collected in a 12-hour time window, then maintained at 25°C. Newly hatched larvae were synchronized and raised at low density (30 larvae/vial) at 18 °C for 4 days, then shifted to 29 °C for 42 hours. BRGCs were dissected from the larvae. Ten BRGCs were put into a 1.5 ml tube, flash frozen in liquid nitrogen, then stored at −80 °C. Three biological replicates were prepared for each genotype. For the quantification of steroidogenic gene expression in *dfr^14^*, brain ring gland complexes were dissected from wandering third instar larvae. Four biological replicates were prepared for control *w^1118^* and *dfr^14^*. RT-qPCR was performed as previously described (Lindberg et al., 2018). The TaqMan probes are as follows: *phm*, Dm01844265_g1; *nvd*, Dm01844265_g1; *sro*, Dm02146256_g1; *dib*, Dm01843084_g1; *sad*, Dm02139319_g1. The measured transcript levels were normalized relative to *Rpl32* values. Statistics was performed using multiple *t*-tests with Holm-Sidak correction in Graphpad Prism 7.

### Flip-out clones

Cell clones were induced as previously described (Zhao et al., 2015) with minor changes. Female virgins *hs-Flp^122^; UAS-Flp^JD1^/CyO, Act-GFP^JMR1^; Act>stop>Gal4, UAS-GFP^LL6^/TM6b* were crossed with *w^1118^, UAS-dfr-RNAi, UAS-dfr-S*, and *UAS-dfr-L*, respectively. Embryos were collected in a 24-h time window in vials with normal fly food and extra yeast. A 7-10 min heat shock was applied in a 37 °C water bath 24 hours after embryo collection to induce flp-out clones. After clone induction, the vials were placed in a room-temperature water bath for 10 min and then kept at 25 °C. To enable a comparative approach, the specimens of different genotypes incubated with each antibody were analyzed using confocal scanning with identical parameters.

### Ecdysteroid measurements

Ecdysteroid levels were measured with an ELISA kit (20-Hydroxyecdysone Enzyme Immunoassay Kit, ARBOR ASSAYS^TM^) according to the manufacturer’s protocol. Ecdysteroids were extracted followed the protocol in (Danielsen et al., 2016). Briefly, whole animals at the designated time points were homogenized in 0.3 ml methanol by a close fitting pestle, followed by shaking for 4 hours, centrifugation at 14,000 g and collection of the supernatant. The remaining tissues were re-extracted with 0.3 ml methanol and then with 0.3 ml ethanol. The supernatants were pooled and 0.3 ml was evaporated using SCANVAC (Coolsafe^TM^) freeze dryer followed by re-suspension in Assay Buffer (ARBOR ASSAYS^TM^). Absorbance was measured at 450 nm.

### Cell transfections and luciferase assays

Cell transfections and luciferase assays were performed in *Drosophila* Schneider line-2 cells (S2 cells) as previously described (Komura-Kawa et al., 2015) with minor changes. Cells were seeded in 100 μl Schneider’s *Drosophila* medium (GIBCO) in a 96-well plate one day before transfection. Cell transfections were performed using the Effectene Transfection Reagent (Qiagen). Two days after transfection, luciferase assays were carried out using the Dual-Luciferase Reporter Assay System (Promega) following the manufacture’s protocol and analysed with the EnSpire plate reader (PerkinElmer). The *Actin5C-Gal4* plasmid(Komura-Kawa et al., 2015) was used to drive the expression of *UAS-dfr-S*, *UAS-dfr-L, HA-Mld-pUAST* (Uryu et al., 2018) or *UAS-GFP* (as control) together with *pGL3-nvd-Fluc* or *pGL3-spok-Fluc* reporters (Uryu et al., 2018). The Copia Renilla Control plasmid (#38093; Addgene) (Lum et al., 2003) was used for measurements of transfection efficiency. Statistical analysis were performe on log^2^ transformed data with One-Way ANOVA with Tukey correction.

## Supporting information

Video linked to Figure 5-supplementary figure 1C

RNA seq data analyses linked to Figure 4

RNA seq data analyses linked to Figure 4

RNA seq data analyses linked to Figure 4

## Acknowledgements

We want to express our thanks to Bloomington *Drosophila* Stock Center, Vienna *Drosophila* RNAi Center, Kim Furbo Rewitz and Takashi Koyama for fly stocks, W. Johnsson for the *dfr/vvl* cDNA clone, Sarah Certel, Michael O’Connor and Ryusuke Niwa for antibodies and R.N. also for luciferase reporter plasmids.

The authors also acknowledge the technical support from the Imaging Facility at Stockholm University, the Proteomics Biomedicum, Karolinska Institute, Sweden, the Agrisera AB, Vännäs, Sweden and the support from the National Genomics Infrastructure in Stockholm funded by Science for Life Laboratory, the Knut and Alice Wallenberg Foundation and Swedish Research Council, and SNIC/Uppsala Multidiciplinary Center for Advanced Computational Science for assistance with massive parallel sequencing, and access to the UPPMAX computational infrastructure.

This research was supported by the Swedish Cancer Society to Y.E., the Swedish Research Council to Y.E. and by Stockholm University to Y.Z. and Y.E.

## Data Availability

Sequencing data have been deposited in GEO under the accession number GSE149972. All other data generated or analysed during this study are included in the manuscript and supporting files.

