## Supplemental figures and legends for "Stop codon readthrough of a POU transcription factor regulates steroidogenesis and developmental transitions"

**Supplementary material linked to Yunpo et al.**

**Supplementary Table 1-3:** RNA seq data analyses linked to Figure 4 have been uploaded as separate excel flies.

**Video:** A video linked to Figure 5-supplementary figure 1C has been uploaded as a separate file.

**Supplementary figure legends:**

**Figure 1-figure supplement 1. *dfr* mRNA is translated into Dfr-S and Dfr-L.**
(**A-D**) Immunoblots using anti-Dfr-N (A, C and D) and anti-Dfr-C (B) antibodies.
**(A)** Dfr-S and Dfr-L are present in different *Drosophila* species, *OrR*, $w^{1118}$, and *Canton-S*. In addition to the bands with predicted molecular weights (46 kDa and 77 kDa), weak bands with slower migration (54-58 kDa and 100kDa) were observed in these adult extracts (asterixes).
**(B)** Immunoblot with the same extracts as in (A) shows the presence of Dfr-L, but not of Dfr-S, demonstrating the specificity of the Dfr-C antibody.
**(C)** *In vitro* translation of *dfr* cDNA produced several bands, with the strongest one matching the expected migration of Dfr-L.
**(D)** Extracts from embryos reveal only a Dfr-S band.
(**E**) No alternative splicing was observed in the *dfr* gene around the first stop codon. Gel electrophoresis of an RT-PCR product of *dfr* mRNA around the first in-frame stop codon. Only a single band was detected.
(**F**) Sequencing results show no indication of RNA editing around the first stop codon. Upper panel, sequenced with forward primer; lower panel, with reverse primer. Stop codon sequences are boxed.

**Figure 2-figure supplement 1. Dfr-L is present in several larval and adult tissues.**
(**A-H**) Confocal images of *Drosophila* tissues stained with anti-Dfr-C (red) and DAPI (blue), of adult brain (A); crop (B); adult salivary gland (C) with an arrow pointing at the tip cells with prominent Dfr-C staining; female oviduct (D) and germarium (E); late stage embryo (F)

and boxed region in magnified view (F'), with arrow pointing at the embryonic ring gland; L3 wing imaginal disc (G) and gonad of female white prepupa (H). Scale bars 50 µm.

(**I**) Summary of anti-Dfr-N and anti-Dfr-C staining in control *($w^{1118}$)* and *$dfr^{14}$* mutant larval and adult tissues. The fluorescence intensity is represented by -, -/+, +, ++, and +++, from barely detectable to strong.

**Figure 5-figure supplement 1. 3-Dimensional structure of a larval brain-ring gland complex.**

(**A**) Schematic illustration of an L3 larva (upper) and a BRGC, lateral view, anterior to the left. RG, ring gland; BL, brain lobe; VNC, ventral nerve cord; D, dorsal; L, lateral; A, anterior. The ring gland is attached to the brain. It also connects to the spiracles and mouth hooks via the trachea. The bilateral trachea interconnect within the gland.

(**B**) 3-D reconstruction of a larval BRGC, posterior view. *Aug-Gal4>UAS-GFP* marks the corpus allatum (green), anti-Sad the prothoracic gland (red), and DAPI stains DNA (blue).

**(C)** 3D movie of a larval BRGC is uploaded as rich media. The ring gland locates above the brain. The BRGC first rotates 360 degrees along the x axis (horizontal rotation), then rotates 180 degrees along the y axis (vertical rotation). The first image of the movie is in posterior view; the last image is in anterior view (upside down).

**Figure 6-figure supplement 1. Overexpression of Dfr-S inhibits expression of *nvd, phm, dib* and *sad* mRNA, while Dfr-L overexpression**

(**A-D**) Quantification of mRNA in extracts of BRGCs using RT-qPCR after reducing *dfr* mRNA by RNAi or overexpression of *UAS-dfr-S* or *UAS-dfr-L* in larval PG using the *$Phm$-$Gal4^{ts}$* driver. Downregulation of *dfr* or *UAS-dfr-S* overexpression significantly reduced the mRNA levels of *nvd, phm, dib* and *sad,* while overexpression of *UAS-dfr-L* had a comparably weaker inhibitory effect on these target genes, albeit significant for all except *sad.* Statistical analysis was performed with One-Way ANOVA with Tukey correction. *p<0.05, **p<0.01

**Figure 7-figure supplement 1 Dfr regulates the expression of Nvd, Phm and Sad**

(**A, C, E**) Prothoracic glands (PGs) carrying GFP-labelled flp-out clones that express different transgenes (control, *UAS-dfr-RNAi*, *UAS-dfr-S*, and *UAS-dfr-L*). Induction of flp-clones as described in Figure 7. The PGs were stained with anti-Nvd (A), anti-Phm (B) and anti-Sad (C), and shown in magenta (upper panels) or grey (lower panels).

Immunofluorescence of Nvd, Phm and Sad was reduced or totally abolished in *UAS-dfr-RNAi*, *UAS-dfr-S*, and *UAS-dfr-L* clones. Scale bars 25 µM.

(**B, D, F**) Quantification of the relative fluorescence in (A), (C) and (E) respectively. Statistical analysis was performed with One-Way ANOVA with Holm-Sidak correction. *p<0.05; **p<0.01; ***p<0.001; ****p<0.0001.

**Figure 8-figure supplement 1**

**(A)** Amino acid length, molecular weight and amino acid composition of the predicted ORF2 of Dipteran Dfr/Vvl proteins.

**(B-H)** Disorder analysis of Dipteran Dfr/Vvl-L proteins. The intrinsic disorder of Dfr/Vvl-L was calculated by the VSL2 algorithm (http://www.pondr.com/). Schematic representation of ORF1 (red), ORF2 (Yellow) and the DNA-binding domains (DBD, blue) are shown above the disorder graph. The horizontal bold line indicates 0.5 disordered score, above which the amino acid sequence is disordered. Abbreviated species name is indicated below the disorder graphs. The graph for *Drosophila melanogaster* (B) is the same as shown in Figure 8D.

### Supplementary Figures:

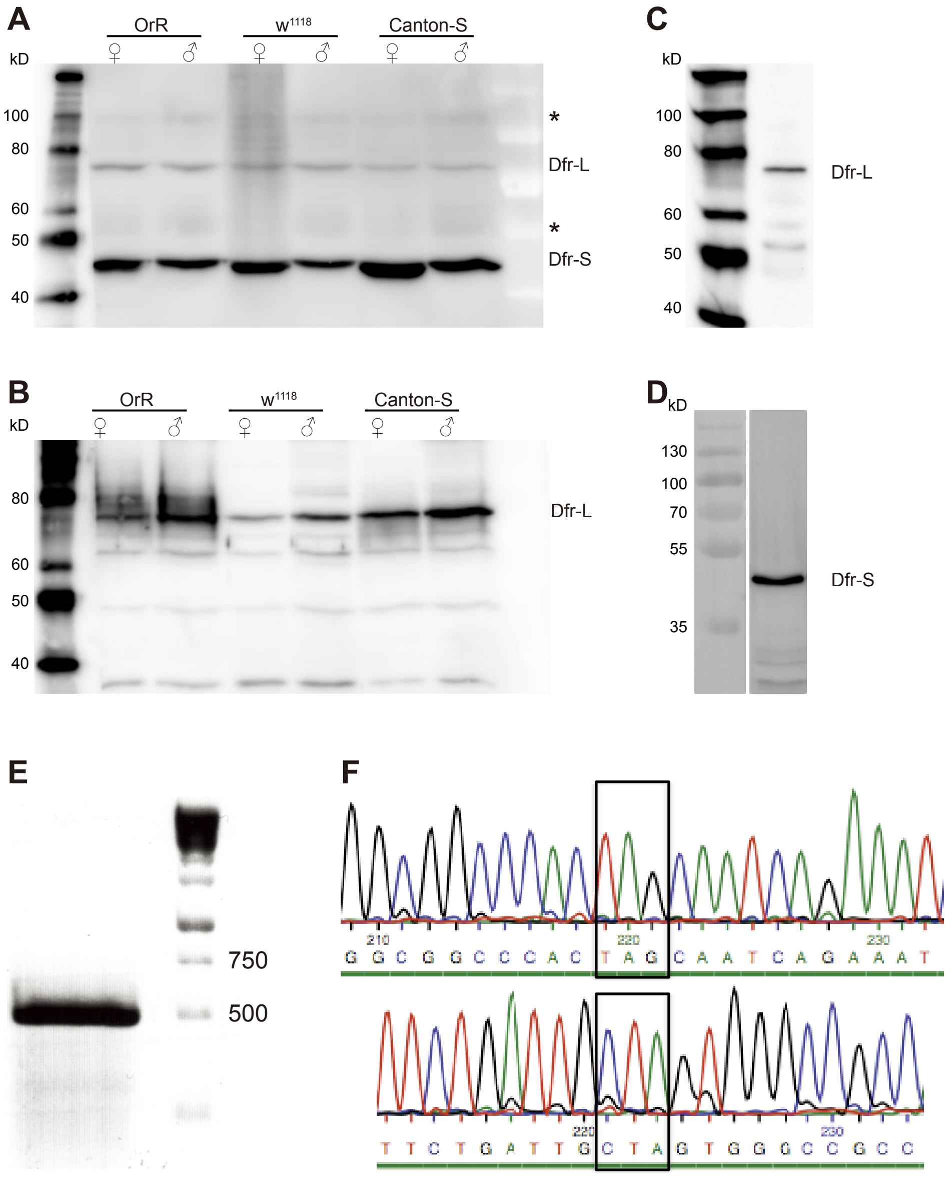


**Figure 1 - figure supplement 1**

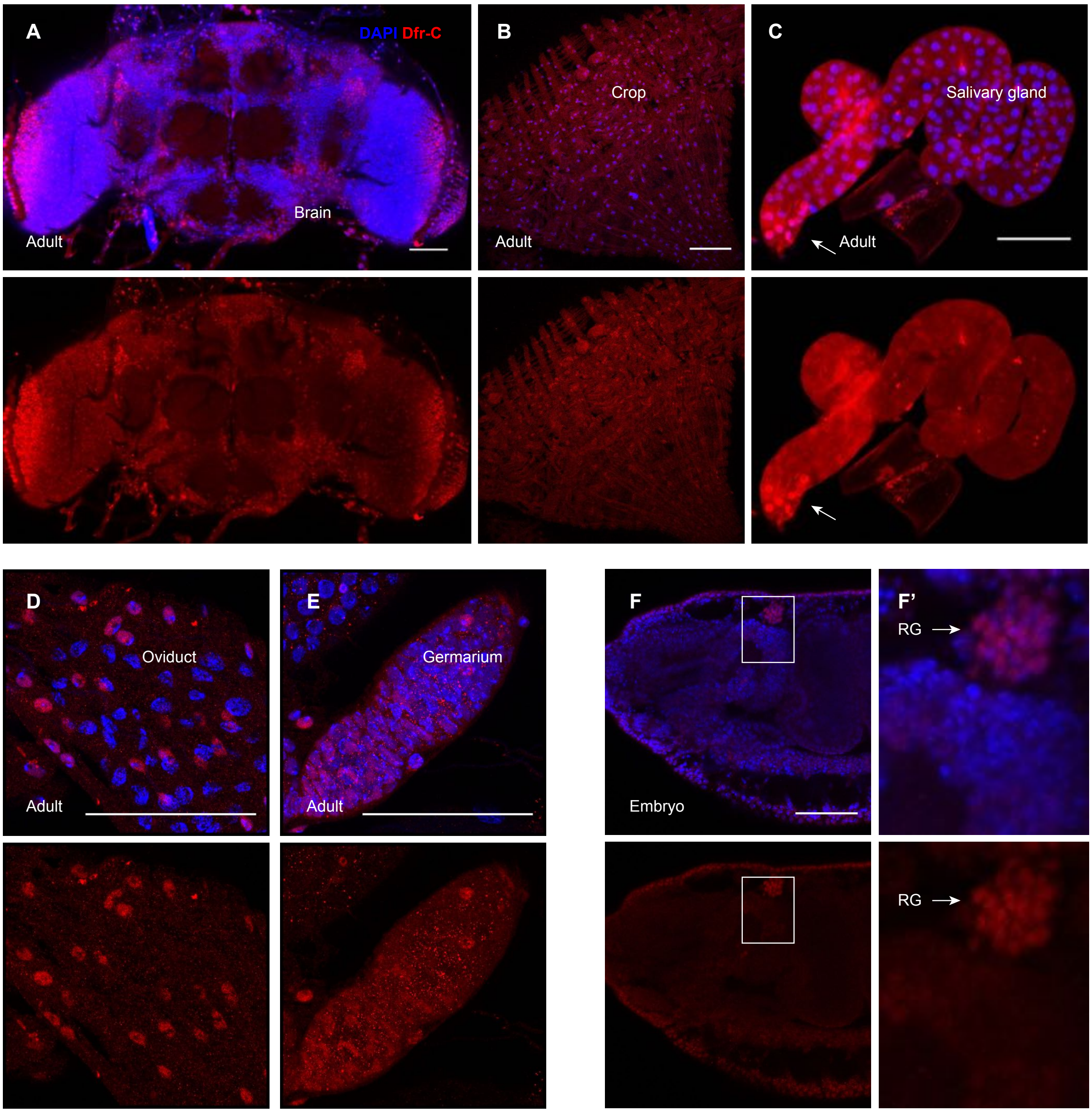


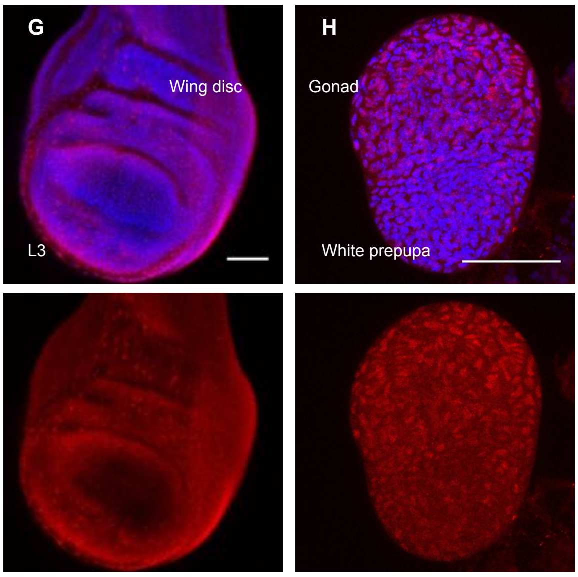


| I | anti-Dfr-N | | anti-Dfr-C | |
|---|---|---|---|---|
| | $w^{1118}$ | $dfr^{14}$ | $w^{1118}$ | $dfr^{14}$ |
| **larva** | | | | |
| L1 prothoracic gland | +++ | +++ | +++ | - |
| L2 prothoracic gland | +++ | +++ | +++ | - |
| L3 prothoracic gland | +++ | +++ | +++ | - |
| L1 brain | ++ | ++ | ++ | - |
| L2 brain | ++ | ++ | ++ | - |
| L3 brain | + | + | + | - |
| trachea | ++ | ++ | ++ | - |
| fat body | + | + | + | - |
| wing disc | + | + | + | - |
| **adult** | | | | |
| ejaculatory duct | +++ | +++ | + | - |
| testis | + | + | + | - |
| ovary | + | + | + | - |
| trachea | ++ | ++ | - | - |
| fat body | ++ | ++ | ++ | - |
| oenocyte | +++ | +++ | ++ | +/- |
| crop | + | + | + | - |
| salivary gland | ++ | ++ | + | - |
| ureter | ++ | ++ | - | - |
| brain | ++ | ++ | ++ | +/- |
| malpighian tubes | + | + | + | - |

**Figure 2 - figure supplement 1**

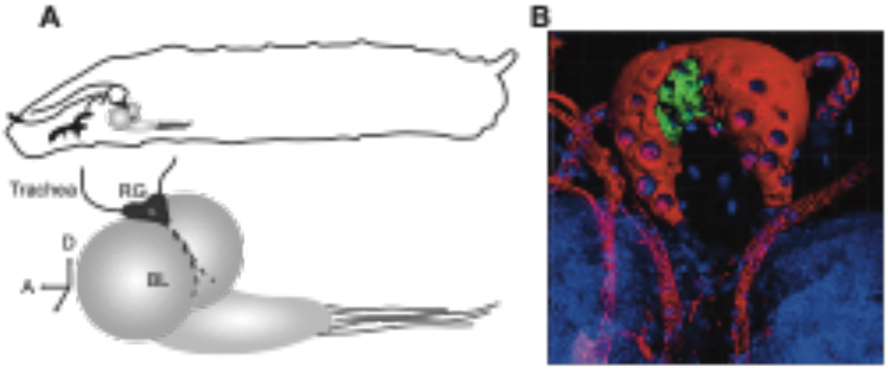


**Figure 5 - figure supplement 1**

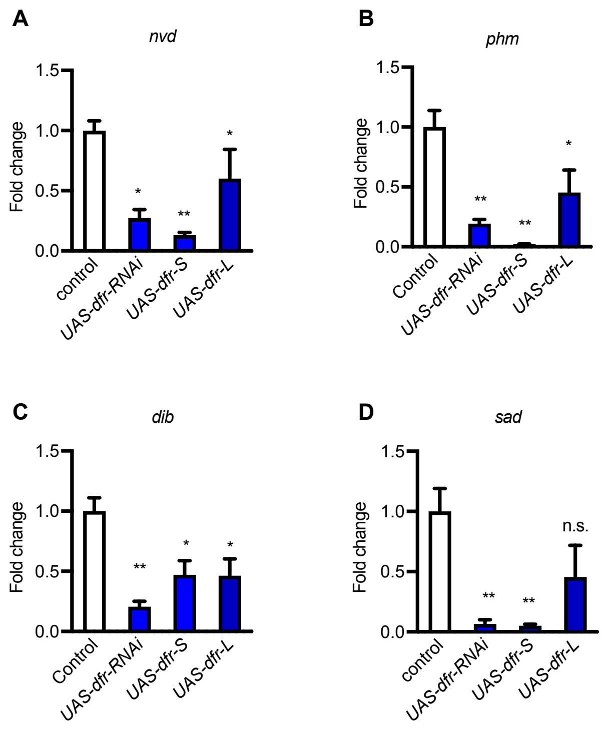


**Figure 6 - figure supplement 1**

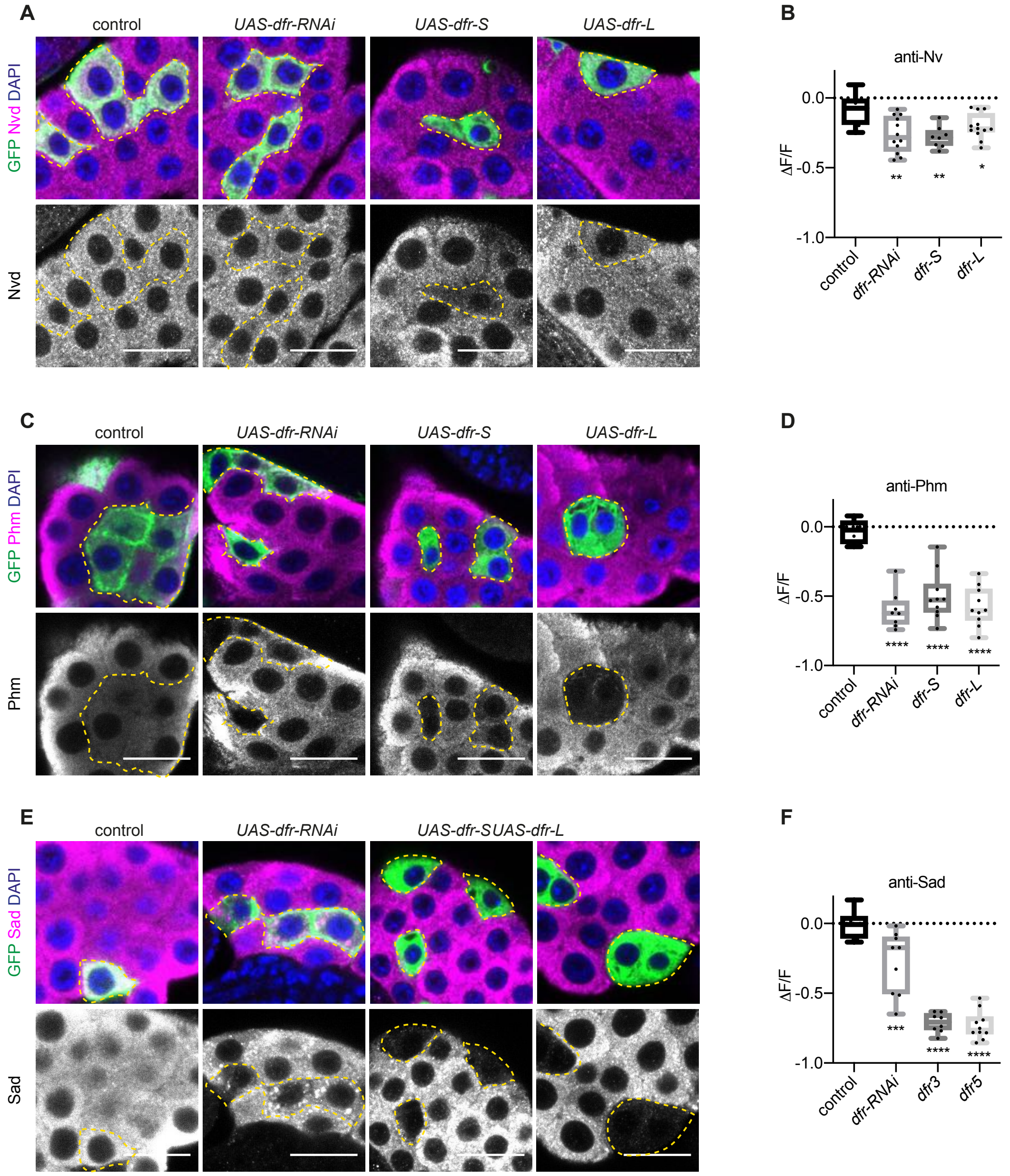


**Figure 7 - figure supplement 1**

A

| Dfr/Vvl protein | ORF2 | ORF2 | ORF2 amino acid composition (%) | | | | | |
|---|---|---|---|---|---|---|---|---|
| Species | lenght (AA) | mol. wt (kDa) | Gln | Ser | His | Ala | Asn | Pro |
| *Drosophila melanogaster* | 286 | 30.9 | 19.9 | 13.3 | 11.5 | 10.5 | 10.5 | 7.0 |
| *Drosophila pseudoobscura* | 313 | 33.9 | 24.0 | 12.5 | 10.9 | 11.8 | 8.0 | 5.8 |
| *Glossinia morsitans* | 269 | 29.0 | 16.4 | 13.0 | 13.8 | 13.0 | 10.4 | 4.8 |
| *Lucilla cuprina* | 261 | 28.3 | 16.5 | 13.8 | 13.8 | 13.0 | 9.6 | 5.4 |
| *Anopheles gambiae* | 199 | 20.1 | 18.6 | 9.0 | 12.6 | 25.6 | 0 | 4.5 |
| *Aedes aegypti* | 126 | 13.1 | 9.5 | 16.7 | 7.1 | 19.0 | 4.0 | 7.1 |
| *Culex quinquefasciatus* | 124 | 13.3 | 10.5 | 14.5 | 11.3 | 12.9 | 4.0 | 8.9 |

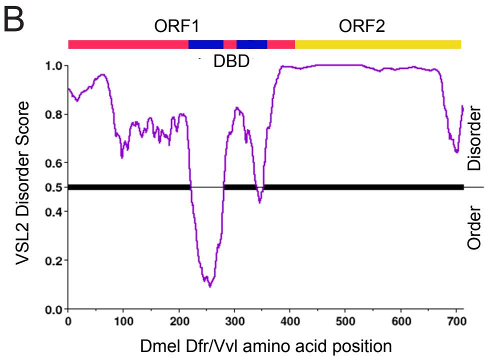


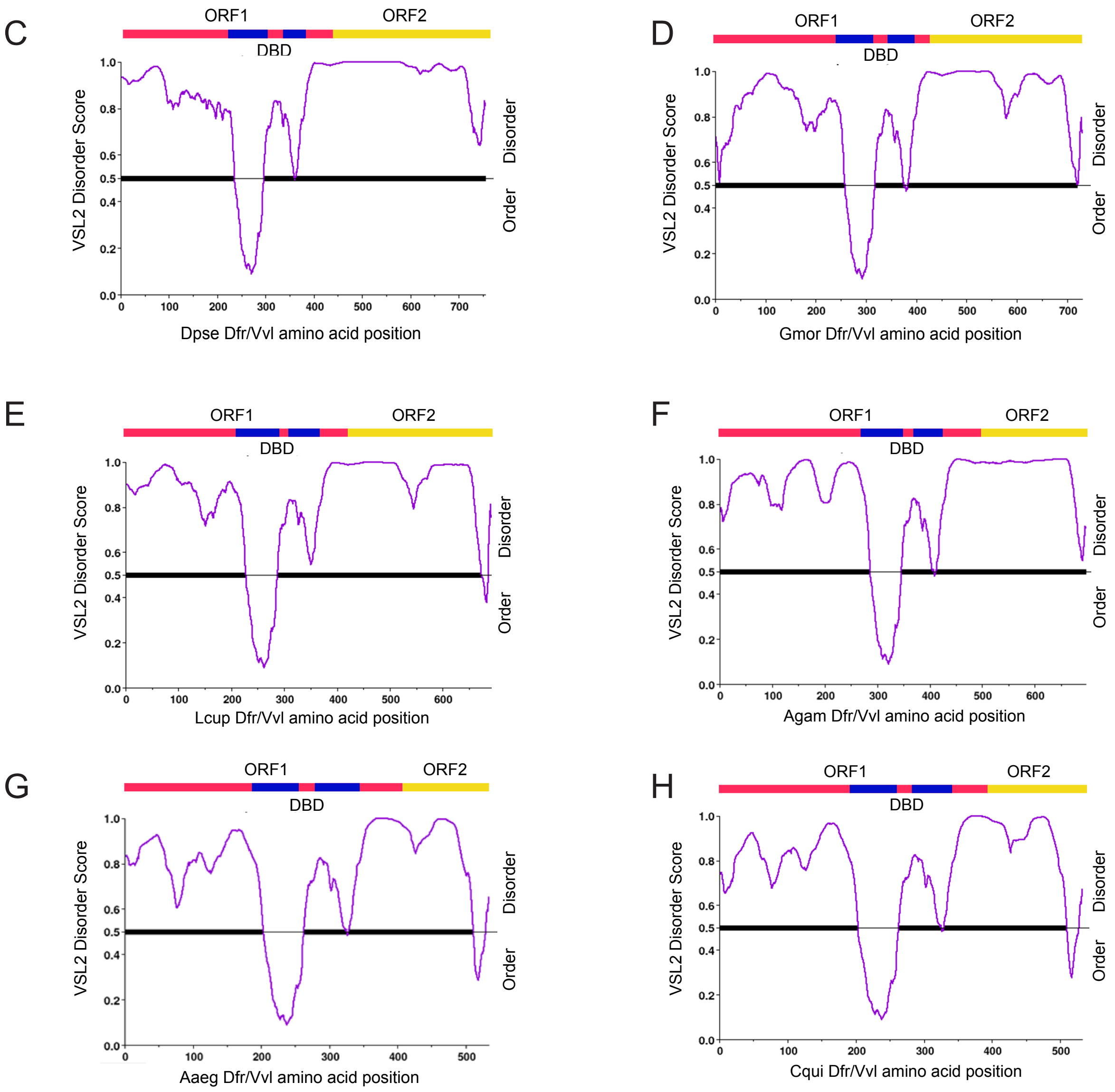


**Figure 8- figure supplement 1**
